# Procentriole microtubules as drivers of centriole reduplication

**DOI:** 10.1101/2020.04.12.038307

**Authors:** Alejandra Vasquez-Limeta, Catherine Sullenberger, Dong Kong, Kimberly Lukasik, Anil Shukla, Jadranka Loncarek

**Affiliations:** Laboratory of Protein Dynamics and Signaling, National Cancer Institute, National Institutes of Health, Frederick, Maryland, 21702, USA

**Keywords:** Centriole, distancing, disengagement, reduplication, Plk1, STORM

## Abstract

Centriole reduplication leads to the formation of supernumerary centrosomes, which promote cellular transformation, invasion and are a hallmark of tumors. A close association between a mother centriole and a procentriole (engagement), established during centriole duplication, intrinsically blocks reduplication. Premature loss of centriole association predisposes centrioles for reduplication and occurs during various types of cell cycle arrests in the presence of high Polo-like kinase 1 activity. Here we use nano-scale imaging and biochemistry to reveal the processes leading to the loss of centriole association and reduplication. We discover that centriole reduplication is driven by events occurring on procentriole microtubule walls. These events are mechanistically different from mitotic centriole separation driven by Pericentrin and Separase but are similar to the physiological process of centriole distancing occurring in unperturbed cycling G2 cells. We propose a concept in which centriole reduplication is a consequence of hijacked and amplified centriole maturation process.

**Highlights:**

- Separase-mediated Pericentrin reorganization is not required for centriole distancing and reduplication in interphase.
- Expression of active Plk1 in S phase leads to centrosomal ultrastructural changes resembling G2 phase.
- Procentrioles without microtubule walls cannot disengage.
- Centriole distancing is intrinsically regulated by the events occurring on procentriole microtubules.

## Introduction

In proliferating vertebrate cells, accurate segregation of genetic material critically depends on two centrosomes, which organize two poles of the mitotic spindle during cell division. Centrosomes are assembled by membrane-less cylindrical structures called centrioles, which, in human cells, are ~240 nm wide and ~500 nm long and built of nine microtubule (MT) triplets (Vorobjev and Chentsov, 1982). On one, proximal, end of the cylinder, centrioles accumulate a highly organized proteinaceous complex called pericentriolar material (PCM), which is the site of numerous centrosomal functions such as microtubule nucleation and centriole duplication. Before mitosis, the PCM of each centrosome expands tin size (Figure 1- figure supplement 1) in a process called centrosome maturation (Kuriyama and Borisy, 1981; Palazzo et al., 2000; Snyder and McIntosh, 1975), increasing the MT nucleating capacity of centrosomes and promoting the formation of two poles of the mitotic spindle. Centrosome maturation is largely driven by the activity of mitotic polo-like kinase 1 (Plk1) (Lane and Nigg, 1996), which peaks in prophase (Golsteyn et al., 1995).

To maintain only two centrosomes per cell, which is vital for the formation of bipolar mitotic spindles and accurate chromosome segregation in mitosis, the formation of centrioles is stringently regulated during the cell cycle. After mitosis, cells inherit two centrioles which both duplicate in the ensuing S phase by initiating a single new centriole (procentriole, PC) ~50 nm away from their MT wall and in an orthogonal orientation (Figure 1- figure supplement 1A; (Anderson and Brenner, 1971; Shukla et al., 2015; Vorobjev and Chentsov, 1982)). Once the PC is initiated, the formation of additional PCs around parental centrioles (mother centrioles, MCs) is inhibited in trans by the PC (Loncarek et al., 2008; Tsou and Stearns, 2006b; Wong and Stearns, 2003). The distance between PC and MC MT walls (here forth referred to as distance) increases during prophase, when the gap between two centrioles can exceed 100 nm (Figure 1- figure supplement 1B; (Shukla et al., 2015; Vorobjev and Chentsov, 1982)). The increase in MC-PC distance coincides with the increase in Plk1 activity, which is also required for centriole distancing (Loncarek et al., 2010; Shukla et al., 2015). Centrioles distanced in prophase do not fully separate before the end of mitosis. After metaphase to anaphase transition, concomitantly with the reduction of PCM, two distanced centrioles finally separate (known as disengagement (Tsou and Stearns, 2006a)) and assemble independent PCM. In addition to its role in distancing, studies show that Plk1 also participates in the final separation of centrioles in late mitosis (Kim et al., 2015; Tsou et al., 2009), possibly by phosphorylating the PCM component Pericentrin, promoting its cleavage by the cysteine protease Separase. Physical separation of centrioles and Pericentrin cleavage at the end of mitosis were suggested to be important for licensing centrioles for duplication in the subsequent cell cycle (Lee and Rhee, 2012; Matsuo et al., 2012; Tsou et al., 2009).

In pathological situations such as tumors, cells accumulate supernumerary centrosomes. Supernumerary centrioles increase the incidence of multipolar mitosis and chromosome miss-segregations, they are a hallmark of aggressive tumors, and mouse models show that they lead to tumorigenesis (Ganem et al., 2009; Nigg and Holland, 2018; Nigg and Raff, 2009; Nigg et al., 2017). In addition, supernumerary centrioles can promote the formation of structurally defective centrioles (Kong et al., 2020). One route to supernumerary centrioles (at least in cell culture conditions) is reduplication. During reduplication, the original PCs prematurely distance from MCs, allowing them to initiate new PCs within the same cell cycle (Loncarek et al., 2008).

Pathological centriole reduplication can occur in cycling or cell cycle arrested cells (Balczon et al., 1995; Mullee and Morrison, 2016; Saladino et al., 2009) which maintain higher than normal Plk1 activity (Loncarek et al., 2010). During centriole reduplication, Plk1 specifically promotes the first step of centriole distancing, while it does not seem to influence the subsequent step of centriole initiation (Loncarek et al., 2008). Although centriole distancing is a critical event leading to reduplication, this process is yet to be understood on a molecular level.

One logical assumption is that, in analogy with processes in cycling cells, Plk1 induces mitotic-like expansion of PCM, pushing PCs further from MCs. The same concept could explain physiological centriole distancing during prophase. Nevertheless, previous microscopy studies of reduplicating centrioles revealed no mitotic-like expansion of PCM component ɤ-tubulin (Shukla et al., 2015). However, optical resolution achieved in these experiments may have not been enough to detect its subtle rearrangements and only a limited number of PCM components were evaluated. In addition, Plk1-driven centriole distancing and reduplication is accompanied by the association of PCM components with PCs, which is called centriole maturation (N.B. different from pre-mitotic centrosome maturation). This observation prompted us to hypothesize that structuring of PCM around PCs during their maturation could be the major driving force for centriole distancing (Shukla et al., 2015). However, this hypothesis has not been directly tested. Finally, it has not been explored whether a Plk1/Separase/Pericentrin-dependent mechanism plays any role in the context of centriole reduplication in interphase.

In this work, we combined nano-scale imaging and biochemical methods to dissect the specific role of Pericentrin, Separase, PCM expansion and PC maturation in centriole distancing and reduplication.

## Results

### Centriole distancing and reduplication in interphase occurs in the absence of Pericentrin remodeling

To study centriole distancing and reduplication, we employed a well-established experimental model (Shukla et al., 2015): HeLa cells were synchronized by shake off and treated with Hydroxyurea before S phase entry. Such cells enter S phase 9-10 h after shake off, duplicate centrioles once, and remain arrested in S phase. Endogenous Plk1 levels in such cells are low, as is typical for S phase cells (Figure 1A). Centriole distancing and reduplication is then promoted by induction of an active mutant of Plk1 (Plk1TD) by doxycycline. In addition, stable expression of centriolar marker Centrin1-GFP allows a direct read out of centriole duplication and reduplication status. To analyze localization of proteins within centrosomes, we used Stochastic Optica Reconstruction Microscopy (STORM), which allows detection of fluorophores with ~22 nm resolution (Bates et al., 2007; Bowler et al., 2019).

**Figure 1.**
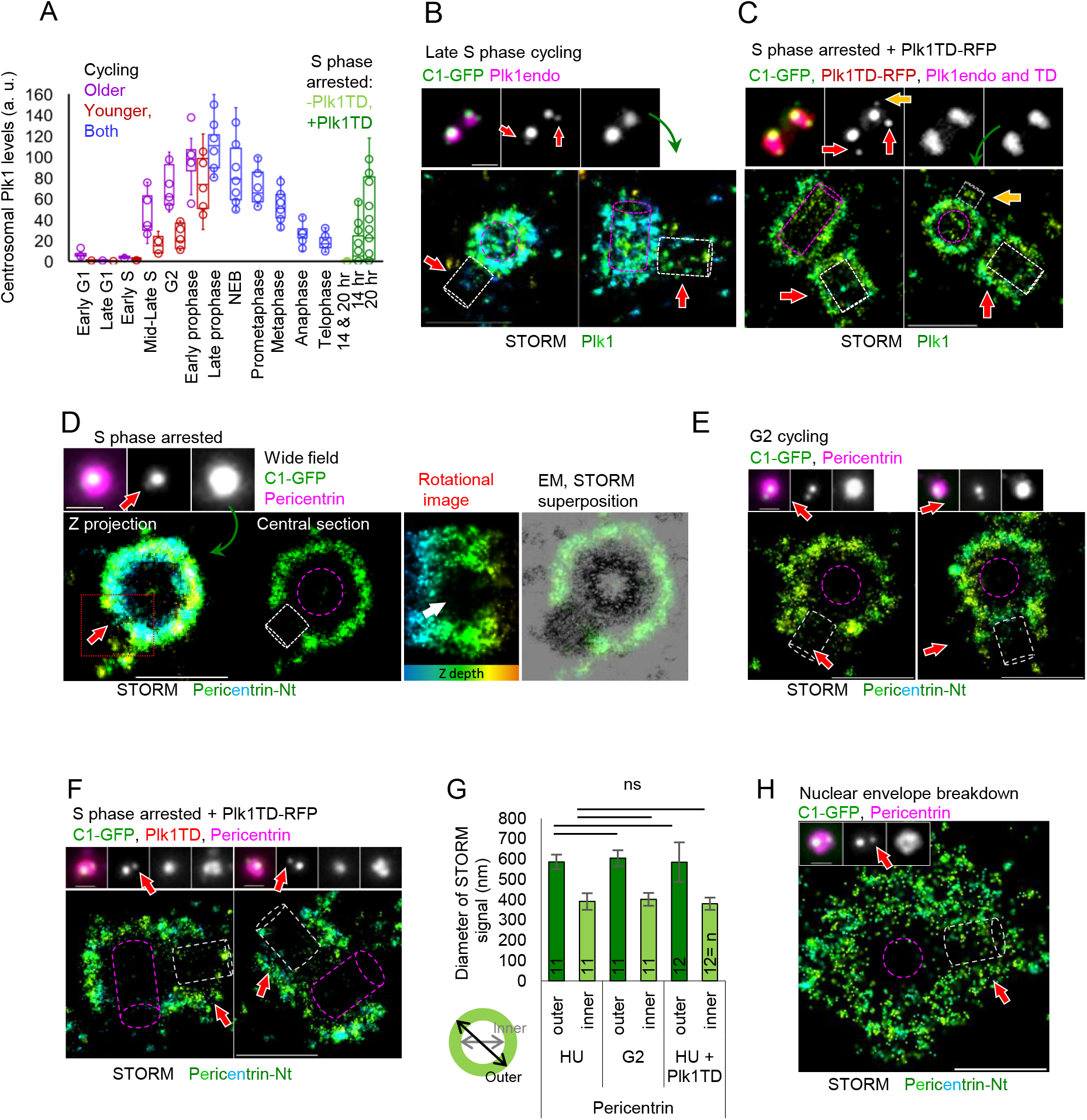
Mother centriole associated Pericentrin is not rearranged during centriole distancing in interphase. (A) Quantification of centrosome-associated endogenous Plk1 and active Plk1 (Plk1TD) levels in cycling and S phase arrested HeLa cells. Centrioles from 10 – 16 cells were analyzed per each point. (B) Correlative widefield/STORM analysis of centrosomal endogenous Plk1 in S phase. Procentrioles are marked by red arrows in all panels. (C) Correlative widefield/STORM analysis of centrosomal Plk1TD in S phase arrested HeLa cells. Yellow arrows mark a procentriole formed in reduplication. (D) Correlative widefield/STORM analysis of centrosomal Pericentrin in S phase arrested HeLa cells. Rotational image of the portion of the STORM signal outlined in red illustrates the gap in Pericentrin signal occupied by the procentriole (white arrow). A superposition of an electronmicrograph of another S phase centrosome with STORM signal, to illustrate the position of centrioles with respect to the typical Pericentrin signal. Electronmicrograph was scaled to match the STORM image. (E) Correlative widefield/STORM analysis of centrosomal Pericentrin in late G2 phase of cycling HeLa cells. (F) Correlative widefield/STORM analysis of centrosomal Pericentrin in S phase arrested HeLa cells expressing Plk1TD. (G) Quantification of STORM signals. Inner and outer diameters were measured as shown in the cartoon. Histograms show an average ± SD from three independent experiments. The sample size (n) is shown on the graphs. ns: nonsignificant. (H) Correlative widefield/STORM analysis of expanded Pericentrin lattice around a centrosome in a cycling HeLa cell undergoing nuclear envelope breakdown. Purple outline: mother centrioles, white outline, procentrioles. Scale bars: 500 nm for STORM images in all panels, 1000 nm for centriole inserts in all panels.

First, we analyzed the sub-centrosomal localization pattern of Plk1 and Pericentrin during centriole distancing. In S phase, centrosome-associated endogenous Plk1 levels are low and increase as cells progress through the cell cycle, reaching highest levels before mitotic entry (Figure 1A). After expression of Plk1TD in S phase arrested cells, the levels of centrosome-associated active Plk1 (Plk1TD) reached the levels of endogenous Plk1 in cycling S, G2 and early prophase cells (Figure 1A). In S phase, both endogenous Plk1 and Plk1TD localized at MCs in a narrow layer surrounding the entire centriole cylinder, enclosing centriole proximal ends (Figure 1B-C and Figure 1- figure supplement 2). Plk1 signal was also present at the MC-PC interface and it was not interrupted by the presence of PCs. Plk1TD, in addition to MCs, abundantly localized to PCs. Initially, Plk1TD was detected at low levels around still orthogonally oriented PCs (Figure 1- figure supplement 2). After several hours, Plk1TD-containing PCs are regularly found at a farther distance and disoriented from MCs (Figure 1C). By the end of our typical experiment (~30 - 32 h after shake off), 20-40% of Plk1TD-positive MCs are usually reduplicated. Characterization of Plk1 dynamics illustrates that our STORM analysis is sensitive enough to detect nano-scale changes of centrosomal proteins within centrosomes.

To understand whether expression of Plk1TD leads to Pericentrin reorganization, we compared Pericentrin localization pattern before and after Plk1TD expression. In S phase cells (cycling or arrested alike), Pericentrin is distributed around MCs with its N-terminus forming a ring of ~600 nm in diameter (Figure 1D and 1G). At the sites of PCs, the ring signal exhibited a circular gap (Figure 1 D; (Mennella et al., 2012)). Cycling G2 cells, characterized by the presence of separated centrosomes and no or minimal DNA condensation, showed similar Pericentrin localization pattern around MCs (Figure 1 E). In addition to MCs, some Pericentrin signal in G2 cells can also be detected in association with PCs (Figure 1E). After Plk1TD expression, centrosomes of S phase cells showed no differences in the pattern of MC-associated Pericentrin, which remained localized in a toroid of similar dimensions as without Plk1TD expression (Figure 1F-G). It is noteworthy that Pericentrin rearrangement during centrosome maturation was readily detectable in cycling cells in our STORM experiments (Figure 1 H). In agreement with a positive effect of Plk1 on PC maturation (Kong et al., 2014; Loncarek et al., 2010; Shukla et al., 2015), expression of Plk1TD induced accumulation of Pericentrin around PCs (Figure 1F). Thus, we concluded that reorganization of MC-associated Pericentrin, as observed during centrosome maturation or mitotic exit, does not occur during centriole distancing and reduplication in interphase. To further test this conclusion, we performed a biochemical analysis of Pericentrin to determine if Separase-dependent cleavage of Pericentrin occurred in S phase cells expressing Plk1TD and additionally explored the role of Separase on centriole distancing and reduplication in S phase.

### Centriole disengagement in interphase does not require Separase

Immunoblotting of Pericentrin showed no Separase-dependent cleavage of Pericentrin in the population of S phase cells expressing Plk1TD, although Pericentrin cleavage was consistently detected in mitotic cells (Figure 2A; (Lee and Rhee, 2012; Matsuo et al., 2012)). Pericentrin cleavage in S phase was not detected even after transient expression of HA-Separase (Figure 2A), although a fraction of centrosome-associated Separase was readily detectable after live-cell permeabilization and removal of over-expressed cytosolic fraction (Figure 2B). In our hands, immunolabeling showed no association of endogenous Separase with S phase or G2 centrosomes, although it was detected on mitotic centrioles, in agreement with previous reports (Agircan and Schiebel, 2014; Chestukhin et al., 2003). Consistently, Separase overexpression had no effect on the dynamics of centriole distancing and reduplication, even after its co-expression with active Plk1 (Figure 2C-E).

**Figure 2.**
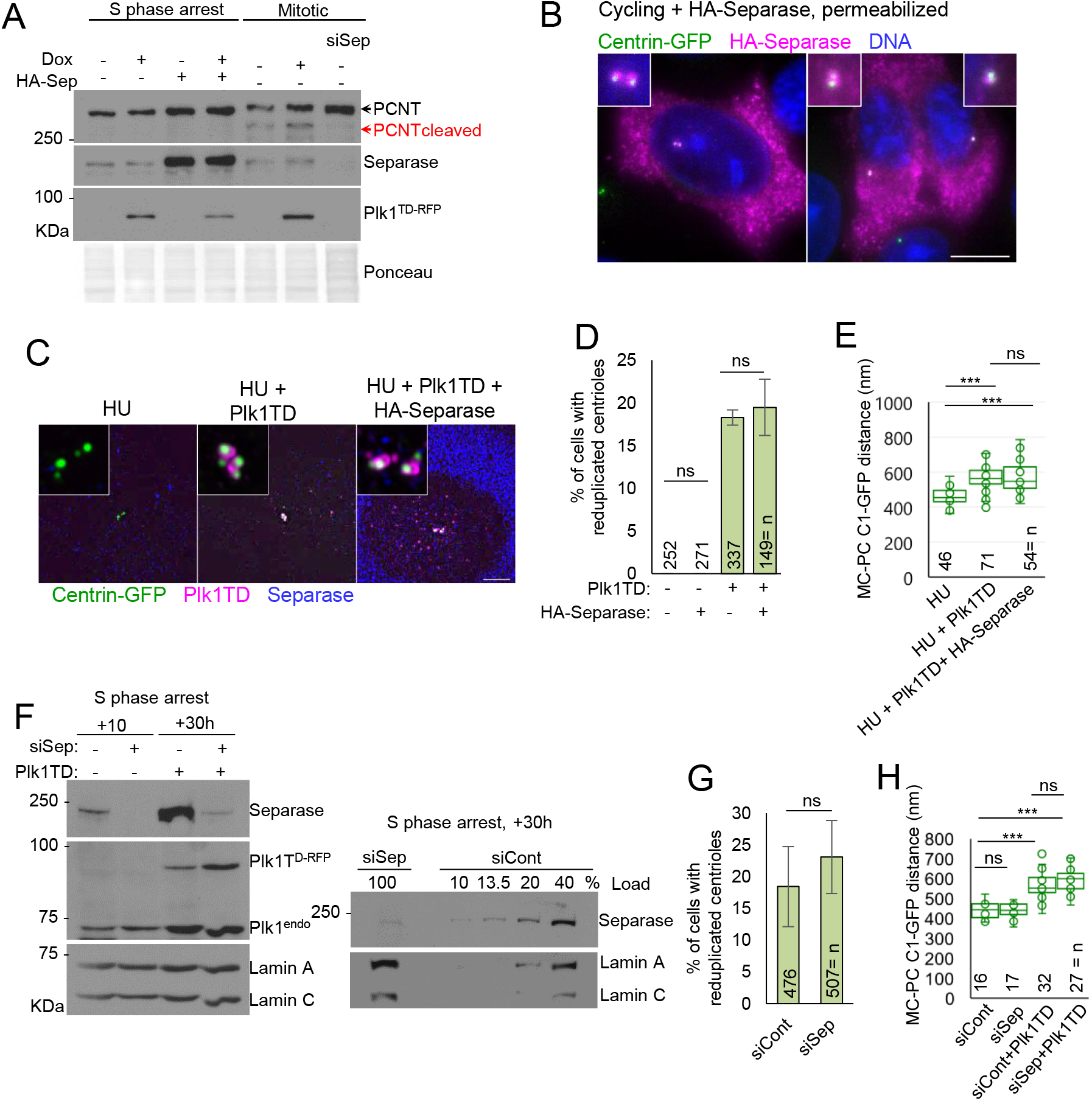
Separase and proteolytic cleavage of Pericentrin do not regulate centriole distancing and reduplication in interphase. (A) Immunoblotting of Pericentrin, Separase and Plk1TD in S phase arrested and cycling mitotic HeLa cells. Active Plk1 (Plk1TD) was induced in some samples by Doxycycline (Dox). Mitotic cells depleted for Separase by siRNA were used as a control. HA-Separase was expressed to increase its levels/activity, and potentially, Pericentrin cleavage in S phase arrested cells. (B) The analysis of HA-Separase localization after expression in cycling cells. Cells were permeabilized before fixation and immunolabeled with anti-Separase antibody. (C) Examples of centrioles in S phase arrested cells by Hydroxyurea (HU) after expression of Plk1TD and HA-Separase. Plk1 and Separase were immunolabeled. (D) Quantification of cells with reduplicated centrioles in S phase arrested, Plk1TD-positive control and HA-Separase-expressing cells. Cells with more than 4 C1-GFP signals were quantified as reduplicated. (E) Quantification of the distance between mother centriole (MC) and procentriole (PC) C1-GFP signals in S phase arrested cells expressing Plk1TD and HA-Separase. (F) Left: Immunoblot analysis of Separase and Plk1 levels in cells depleted of Separase (siSep) (please see Figure 2- figure supplement 1A for experimental details). Right: To quantify the level of Separase depletion, different amounts of control (siCont) sample were loaded alongside siSep sample. 100%: 20 μg of total lysate. (G) Quantification of cells with reduplicated centrioles in S phase arrested cells expressing Plk1TD after Separase depletion. Cells with more than 4 C1-GFP signals were quantified as reduplicated. (H) Quantification of the distance between MC and PC C1-GFP signals in S phase arrested Separase-depleted cells expressing Plk1TD. Histograms show an average ± SD from three independent experiments. The sample size (n) is shown on the graphs. ns: nonsignificant; *: P ≤ 0.05; **: P ≤ 0.01; ***: P ≤ 0.001. Scale bar in D: 4 μm.

Previous studies showed that in proliferating cells, Separase removal (Tsou et al., 2009) or the inhibition of Pericentrin cleavage (Kim et al., 2015; Lee and Rhee, 2012; Matsuo et al., 2012) inhibits or at least delays centriole licensing for duplication in the ensuing cell cycle. These effects were attributed to the lack of centriole disengagement. So, we tested whether Separase depletion perturbs the dynamics of PC distancing and reduplication in S phase arrested cells. Cells were transfected with Separase-targeting or non-targeting siRNA, mitotic cells were harvested ~24 h later, and re-plated. Reduplication assay was performed in the second S phase following siRNA transfection. In parallel, some cells were allowed to cycle (for experimental design please see Figure 2- figure supplement 1A). The efficiency of Separase depletion was determined by immunoblotting, immunostaining and by the presence of chromosome segregation defects in cycling cultures (Chestukhin et al., 2003). Immunoblotting revealed that Separase was depleted by ~90% (Figure 2F). Immunofluorescence showed that Separase was largely undetectable in ~90% of G2, mitotic and G1 cells, while the remaining cells still contained Separase signal albeit weaker than that detected in control cells (Figure 2- figure supplement 1C). In agreement, Separase negative mitotic and postmitotic cells showed reduced cleavage of Pericentrin (Figure 2- figure supplement 1B) and exhibited errors in chromosome segregation, evident by the presence of sister cells associated with DNA bridges and tetraploid cells formed by cell fusion (Figure 2- figure supplement 1D). Nevertheless, quantification of Centrin1-GFP signals in S phase arrested cells expressing Plk1TD, collected in parallel, revealed that in both control and Separase-depleted cells, centrioles distanced and reduplicated at a similar rate (Figure 2G and H). Thus, our data indicate that Separase does not mediate centriole distancing and reduplication in interphase.

### Cycling cells lacking Separase can license centrioles for duplication

Our data showed that in S phase arrested cells, Separase does not regulate neither centriole distancing nor reduplication. To better understand how the interphase disengagement process differs from those in mitosis, we depleted Separase in cycling cells and conducted a detailed STORM analysis of centrosomes in postmitotic cells. Cells were transfected with Separase or with control siRNA and fixed ~48 h after transfection when they were in their second cell division (Figure 2- figure supplement 1A). Cells were immunolabeled for centriole initiating factors Cep152, Cep63 (Kim et al., 2013; Wei et al., 2020), SAS-6 (Dammermann et al., 2004; Leidel et al., 2005), Pericentrin and Separase in various combinations, and analyzed by STORM. Separase-depleted cells were identified by the presence of DNA bridges between two sister cells and the presence of tetraploid fused cells (Figure 2- figure supplement 1D and E). In agreement with reduced Pericentrin cleavage observed by immunoblotting (Figure 2- figure supplement 1B), Separase-depleted postmitotic cells had higher levels of centrosome-associated Pericentrin (Figure 2- figure supplement 1D and E) and MCs and PCs were often positioned closer than in control samples (Figure 2- figure supplement 2A - D). Nevertheless, STORM revealed that albeit adjacent, most PCs were distanced and disoriented from MCs. Further, both PCs and MCs were regularly associated with maturation markers such as Cep63 and Cep152 (Figure 2- figure supplement 2A - D). SAS-6 was absent from centrosomes and cytoplasm of postmitotic Separase-depleted cells similar to control G1 cells (Strnad et al., 2007). S phase re-entry was recognized by the presence of centriole initiator SAS-6, which revealed that some sister cells associated with DNA bridges enter S phase asynchronously (Figure 2- figure supplement 2E). Plk4, a kinase that drives PC initiation (Bettencourt-Dias et al., 2005; Habedanck et al., 2005) and accumulates on centrioles in a Cep63- and Cep152-dependent manner, was, in G1 cells, associated with both centrioles, signifying that both were mature (Figure 2- figure supplement 2E).

Based on our current understanding of the centriole intrinsic block to reduplication and centriole assembly pathway, the presence of centriole maturation markers Pericentrin, Cep63, Cep152 and Plk4 on postmitotic centrioles is not consistent with the lack of PC maturation and licensing for duplication. Indeed, at later time points, SAS-6 positive cells duplicated centrioles, although in some fused cells centriole duplication was asynchronous, consistent with previous observations (Tsou et al., 2009). We reason that delayed and asynchronous centriole duplication that can be observed in Separase-depleted cells might be a consequence of perturbations in cell cycle progression that follow mitotic abnormalities and aneuploidy. In this respect, it is notable that Separase depleted postmitotic cells asynchronously reach S phase (Figure 2- figure supplement 2E). Subsequent fusion of such two cells could explain asynchronous centriole duplication in some tetraploid cells. It is also possible that errors in chromatid segregation deprives some G1 cells of genes needed for proper centrosome function. Finally, it is also plausible that Separase depletion in mitosis perturbs centriole initiation nonspecifically by perturbing centrosome composition. For instance, over-abundance of Pericentrin, which is a consequence of Separase depletion, is not physiological for postmitotic human centrioles. It has yet to be determined whether Separase depletion directly affects other centrosomal proteins.

### Cep192 and ɤ-tubulin expand to G2 levels during centriole distancing and reduplication

Phosphorylation of Pericentrin by Plk1 in prophase of cycling cells, promotes centrosomal accumulation of other PCM components (Lee and Rhee, 2011). Thus, we reasoned that although our microscopy did not detect Pericentrin remodeling during Plk1-induced centriole distancing and reduplication, its phosphorylation might have occurred but remained undetected by STORM. Such phosphorylation could affect the localization of other PCM components, such as Cep192 (SDP-2 in *C. elegans* and *Drosophila)* and ɤ-tubulin, which are dependent on Pericentrin phosphorylation status for their centrosomal recruitment (Lee and Rhee, 2011). Cep192 is an important PCM recruiting factor during centrosome maturation, depletion of which reduces centrosomal levels of ɤ-tubulin, Pericentrin and CDK5Rap2 (Gomez-Ferreria et al., 2007; Haren et al., 2009; Joukov et al., 2014; O’Rourke et al., 2014). Finally, centrosomal Cep192 levels sharply increase in prophase in a Plk1-dependent manner (Figure 3, Figure 3- figure supplement 1A; (Gomez-Ferreria et al., 2007)), coinciding with the distancing of PCs in cycling cells. So, we next tested whether G2 levels of active Plk1 in S phase cells changes localization of two key mitotic PCM components, Cep192 and ɤ-tubulin.

**Figure 3.**
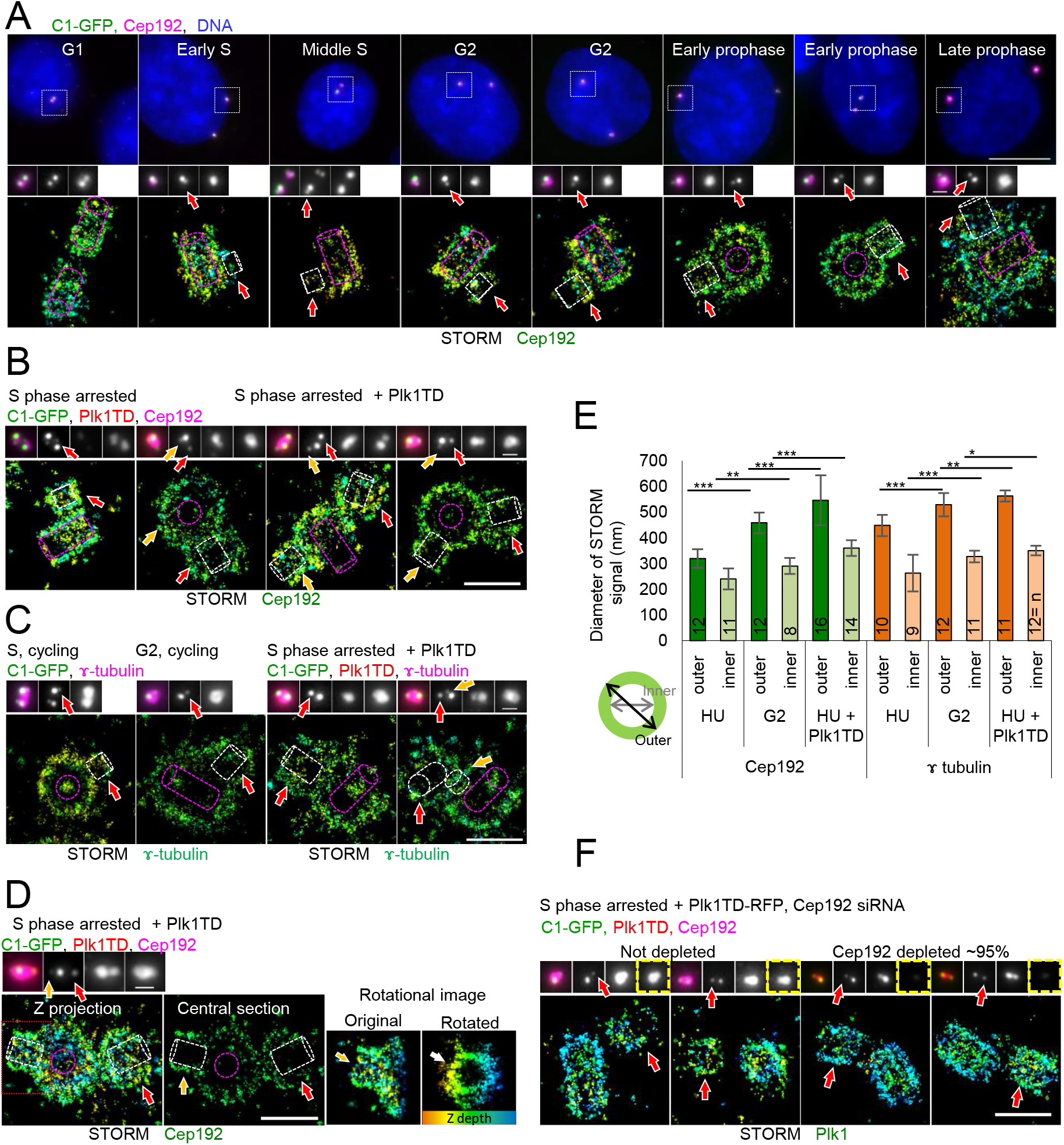
Localization of Cep192 and ɤ-tubulin during centriole distancing and reduplication. Correlative widefield/STORM analysis is shown in all panels. Red arrows point to original procentrioles, yellow arrows point to procentrioles formed during reduplication. (A) Cep192 centrosomal localization during various cell cycle phases in cycling HeLa cells. Centrioles outlined in widefield images are in inserts and were analyzed by STORM. (B) Cep192 centrosomal localization in cells arrested in S phase expressing active Plk1 (Plk1TD). (C) ɤ-tubulin centrosomal localization in cycling and S phase arrested cells expressing Plk1TD. (D) Cep192 centrosomal localization in a cell arrested in S phase expressing active Plk1 (Plk1TD). Rotational image of the portion of the STORM signal outlined in red illustrates the gap in Cep192 STORM signal occupied by procentriole formed during reduplication (white arrow). (E) Quantification of Cep192 and ɤ-tubulin STORM signals. Inner and outer diameters were measured as shown on the cartoon. Histograms show an average ± SD from three independent experiments. The sample size (n) is shown on the graphs. ns: nonsignificant; *: P ≤ 0.05; **: P ≤ 0.01; ***: P ≤ 0.001. (F) Plk1TD centrosomal localization in S phase arrested cells after depletion of Cep192 by siRNA. Plk1TD strongly associates with all centrioles and procentrioles regardless of the presence or absence of Cep192 (yellow squares) and PCs are disoriented and distanced from MCs. Purple outlines: mother centrioles, white outlines: procentrioles. Scale bars: widefield in A: 10 μm. STORM: 0.5 μm, centriole inserts: 1 μm

In S phase of cycling or arrested cells, Cep192 was detected in a thin layer around MCs, forming a toroid of discrete dimensions (Figure 3A). Changes in localization of MC-associated Cep192 was detected already in G2, before obvious signs of DNA condensation. In G2, Cep192 appeared in an additional layer of larger diameter before the signal, by late prophase, became less structured (Figure 3A). In late prophase, toroid-like organization was no longer obvious and Cep192 was localized at a greater distance from centrioles, consistent with its role in centrosome maturation. In cycling cells, starting from G2, Cep192 was also associated with PCs (Figure 3A). Unlike Pericentrin, Plk1TD expression in S phase cells significantly changed the localization of Cep192 (Figure 3B and E). The diameter of MC-associated Cep192 increased in comparison to Plk1TD-negative cells (Figure 3B) or to cycling prophase cells treated with Plk1 inhibitor (Figure 3- figure supplement 1A) and its diameter and pattern resembled G2 centrosomes of cycling cells (Figure 3B). Changes in Cep192 were accompanied by its increased centrosomal levels (Figure 3- figure supplement 1B), mimicking the dynamics of these proteins in cycling G2/prophase cells (Figure 3A). In addition to MCs, Plk1TD expression promoted Cep192 association with PCs, forming larger toroids than found on control cycling PCs in G2 (Figure 3B). ɤ-tubulin localized alongside MC MTs, in the lumen of MCs (Figure 3C; (Fuller et al., 1995)) and with PCs during interphase. The diameter of MC-associated ɤ-tubulin signal increased in G2, as well as after Plk1TD expression in S phase (Figure 3C and E). Plk1TD expression resulted in the PC-associated ɤ-tubulin signal of a larger diameter than in S phase cells. Expansion of Cep192 and ɤ-tubulin in response to Plk1TD expression was permanent, but it did not perturb the formation of new PCs, which formed in the vicinity of MCs and were surrounded by such expanded PCM (Figure 3D).

In addition, inner PCM proteins Cep63 and Cep152 were also included in analysis, as they localize at the MC-PC interface and are critical for PC initiation (Kim et al., 2013). STORM analysis of MC-associated Cep63 and Cep152 revealed ring-like signals (Figure 3- figure supplement 1C). Cep63 localized closer to MCs, while Cep152 formed a ring with a larger diameter, consistent with previous studies (Kim et al., 2019b; Lawo et al., 2012; Mennella et al., 2012; Sonnen et al., 2012). Interestingly, in cycling cells, Cep63 and Cep152 signals were also already associated with the proximal ends of PCs during G2/prophase (Figure 3- figure supplement 1C). Their PC levels were variable, and PCs associated with more Cep63 and Cep152 were found further from MCs (not shown). Unlike on MCs, PC-associated Cep63 and Cep152 signal was not detectable or barely detectable in mitosis and became again detectable in early G1 (Figure 2- figure supplement 2A). Plk1TD expression resulted in no change of MC-associated Cep63 and Cep152, but it promoted their robust accumulation around PCs (Figure 3- figure supplement 1C and D).

In summary, our STORM analysis revealed that, in some respects, expression of Plk1TD in S phase induces centrosomal changes which are characteristic for G2/prophase cells. This opened a possibility that expansion of MC-associated Cep192 and ɤ-tubulin drives PC distancing. Importantly, however, our STORM analysis also revealed that PCs of cycling cells associate with a substantial amount of PCM components already in G2 phase, considerably reinforcing our hypothesis that PC distancing is promoted by PC maturation. Thus, we designed a set of experiments to clarify the role of MC PCM expansion and PC maturation in centriole distancing.

### Mother centriole PCM expansion is not critical for centriole distancing and reduplication

To prevent accumulation of PCM components, we depleted Cep192 in S phase arrested cells and subsequently expressed Plk1TD. Quantification of residual centrosomal Cep192 signal in the cell population showed that Cep192 was, in some cells, depleted to ~95% (Figure 3F). Nevertheless, in Cep192-depleted cells, Plk1TD localized to both centrioles and PCs were found at larger distances than in control cells and were disoriented. Thus, lack of Cep192 accumulation around MCs did not prevent PC association of Plk1TD nor PC distancing. Removal of Cep192 and Spd-2 has repeatedly been shown to reduce the accumulation of other PCM components (Gomez-Ferreria et al., 2007; Haren et al., 2009; O’Rourke et al., 2014; Pelletier et al., 2004). Based on this and on the behavior of other analyzed PCM components, we concluded that the expansion of MC’s PCM during S phase is likely not critical for PC distancing.

### Procentrioles lacking microtubules are stable and maintain centriole reduplication block

In ensuing experiments, we explored the contribution of PC maturation on distancing. The ability of PCs to mature and accumulate PCM components requires the presence of MTs (Atorino et al., 2020; Wang et al., 2017). Thus, we decided to generate PCs that lack MT walls and explore how they behave in the presence of high levels of active Plk1. To generate such PCs, we used Nocodazole, an inhibitor of microtubule polymerization, and the strategy described in Figure 4- figure supplement 1. Briefly, mitotic cells were collected by shake off and re-plated on coverslips. Nocodazole was added while cells were still in G1. Hydroxyurea was added to arrest cells in S phase. In Nocodazole-treated cultures as well as in controls, cells reached S phase and duplicated centrioles ~9-10 h after shake off, as determined by the appearance of a weak C1-GFP signal in the vicinity of each MC-associated C1-GFP signal, meaning that cytosolic microtubules are not required for PC initiation (Figure 4A). To induce distancing and reduplication, doxycycline was added to the culture, as required.

**Figure 4.**
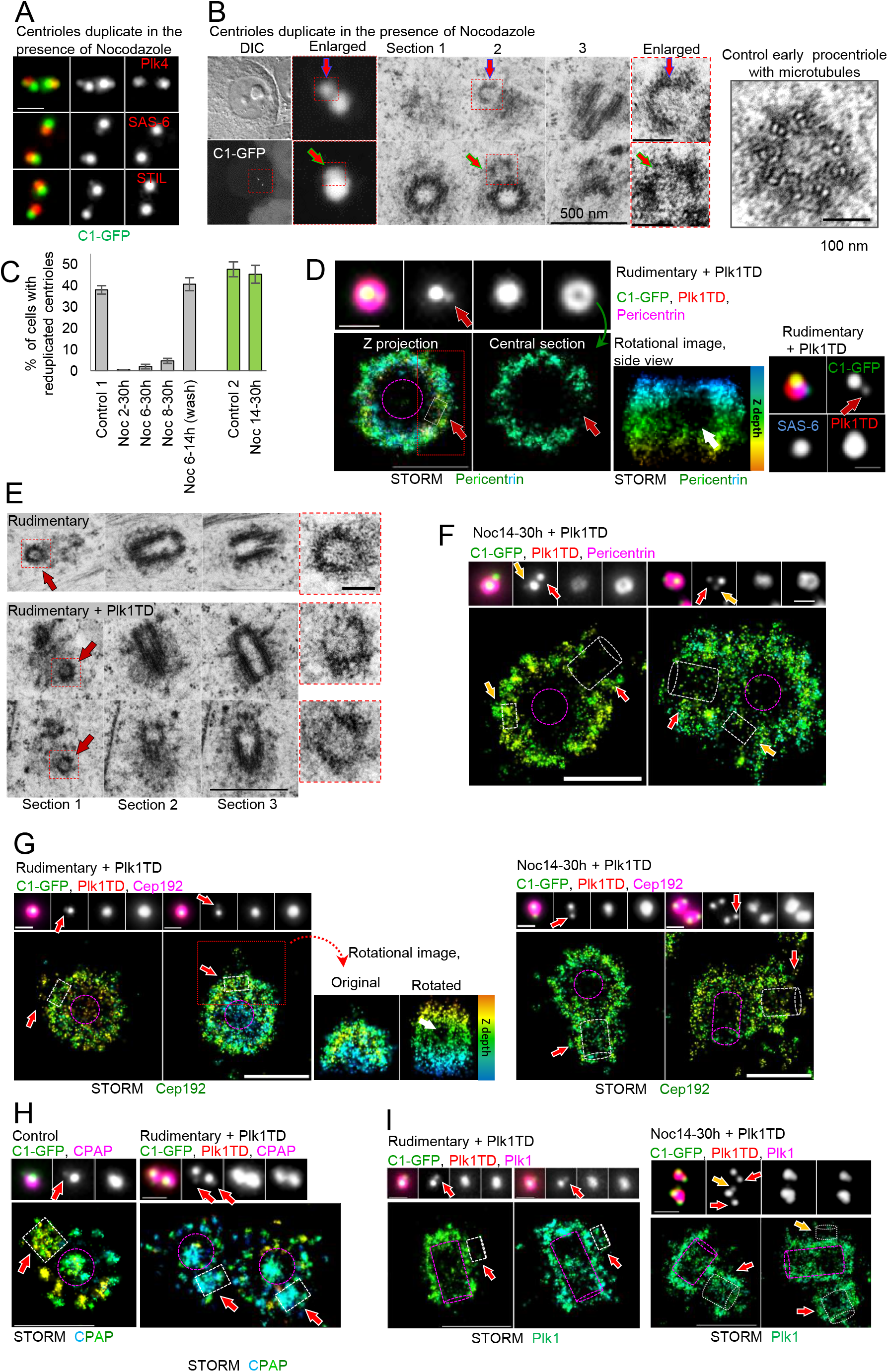
Procentrioles lacking microtubules are stable and maintain centriole reduplication block. For details of experimental strategy used to generate rudimentary procentrioles (RPCs) lacking microtubule walls, please see Figure 4- figure supplement 1. (A) Immunofluorescence analysis of cartwheel proteins STIL, SAS-6 and Plk4 on centrosomes containing RPCs. The signals correlate with weaker C1-GFP signals, indicating procentrioles. Brighter C1-GFP signals label mother centrioles. (B) Correlative live/electron microscopy analysis of RPCs. Red arrows point to procentrioles in all panels. RPCs do not contain microtubule walls and have poorly or no discernable cartwheel. Far right panel: An early control procentriole with assembled microtubule doublets in its wall. Microtubules enclose centriole lumen which contains cartwheel components, centrally positioned hub and radially organized spokes. (C) Comparative widefield/STORM analysis of centrosomal Pericentrin shows a close association of RPC with mother centriole in the presence of high levels of active Plk1 (Plk1TD). Rotational image of the portion of the STORM signal outlined in red illustrates the gap (white arrow) in Pericentrin signal occupied by RPC (red arrow). (D) Quantification of centriole reduplication under various treatment conditions. Cells with more than 4 C1-GFP signals were quantified as reduplicated. (E) Correlative live/electron microcopy analyses of RPCs after ~20h of Plk1TD expression. Three 80 nm-thick serial sections are shown for each centrosome. (F) Localization of Pericentrin on centrosomes containing control procentrioles in S phase arrested cells expressing Plk1TD in the presence of Nocodazole. Red arrows point to original procentrioles. Yellow arrows point to procentrioles formed during reduplication in the presence of Nocodazole. (G) Distribution of Cep192 on centrosomes containing RPCs (left) or control procentrioles (right) in S phase arrested cells expressing Plk1TD in Nocodazole. Rotational image of the portion of the STORM signal outlined in red illustrates the gap (white arrow) in Cep192 STORM signal occupied by the RPC (red arrow). (H) Left: Localization of centrosomal CPAP in control S phase cell by STORM. CPAP localizes in the PCM and in the lumen of mother centrioles and in the lumen of procentrioles. Right: CPAP localization on centrosomes containing RPCs and expressing Plk1TD. CPAP localizes in the PCM and in the lumen of mother centrioles and abundantly colocalizes with RPCs (I) Left: Plk1TD localization within centrosomes containing RPCs. Plk1 does not accumulate to the site of RPCs. Right: Plk1 associates with control procentrioles in the presence of Nocodazole. Yellow arrow points to procentriole formed during reduplication in the presence of Nocodazole. Purple outlines: mother centrioles, white outlines: procentrioles. Scale bars: electron microscopy in B and D: 0.5 μm and 0.1 μm in inserts. STORM in all panels: 500 nm. Widefield centrosome inserts: 1 μm.

The analysis of centrosomes containing PCs formed in the presence of Nocodazole showed that dimmer C1-GFP signals colocalized with cartwheel proteins Plk4, SAS-6 and STIL (Figure 4A). Their characterization by correlative light and electron microscopy showed that weaker C1-GFP signals colocalize with ~90 nm long structures that contained electron transparent ~70 nm-wide lumen surrounded by an unstructured electron denser material (Figure 4B). These structures, which we named rudimentary procentrioles (RPCs) lacked MTs and resembled generative discs previously described in paramecium (Dippell, 1968). A cartwheel, a nine-fold symmetrical structure which is present in control PCs (Figure 4A, right panel; (Anderson and Brenner, 1971; Hirono, 2014; Vorobjev and Chentsov, 1982) was not clearly discernable in RPCs. Nevertheless, such RPCs could implement a sustained block to MC reduplication and were stable during prolonged S phase arrest.

### Rudimentary procentrioles do not distance in the presence of active Plk1

To test how RPCs respond to active Plk1, we induced Plk1TD expression in S phase arrested cells containing RPCs and analyzed centrosomes after ~20 h of sustained Plk1TD expression (~30 h after shake off). By that time, most of the control PCs had already distanced and ~40% of MCs had already reduplicated (Figure 4C). In sharp contrast, RPCs remained closely associated with MCs, and their reduplication was inhibited (Figure 4C and D). Even in cells with the highest levels of Plk1TD overexpression, RPCs remained embedded within Pericentrin and cartwheel protein SAS-6, which is ultimately lost from PCs after distancing, was continuously associated with RPCs (Figure 4D). Correlative live and electron analysis confirmed that after Plk1TD expression, RPCs remained intact and adjacent to MCs even after a 20 h-long exposure to Plk1TD (Figure 4E, n= 8). To eliminate the possibility that Nocodazole treatment nonspecifically inhibits PC distancing, we expressed Plk1TD in S phase arrested cells and treated them with Nocodazole after MCs had duplicated and PCs had already formed MT walls (Figure 4- figure supplement 1). In such cells, Pericentrin signal accumulated on PCs, PCs distanced, and MCs reduplicated (Figure 4F). Quantification of cells with reduplicated centrioles showed that in such samples MCs reduplicated with similar dynamics as in control Nocodazole-untreated samples (Figure 4C, green bars). This experiment demonstrated that the centriole cycle continues in the presence of Nocodazole and in the absence of cytosolic MTs. The analysis of Cep192 showed that centrosomes containing RPCs could still expand Cep192 to the dimensions found in control centrosomes undergoing centriole distancing and reduplication (Figure 4G). The site of RPCs was manifested as a slight reduction of Cep192 signal in otherwise expanded Cep192 signal (Figure 4G, rotated image). Control experiment showed that Cep192 could accumulate on control PCs in S phase Nocodazole-treated cells expressing Plk1TD (Figure 4G, right panel).

Previous studies have demonstrated that the centriole protein CPAP (Kohlmaier et al., 2009; Schmidt et al., 2009; Tang et al., 2009) is critical for Plk1-dependent centriole distancing (Shukla et al., 2015), so we determined whether or not RPCs are able to recruit CPAP. On centrosomes of control S phase cells, CPAP signal localizes in the lumen and in the PCM of MCs and in the cartwheel of PCs (Figure 4H; (Kleylein-Sohn et al., 2007)). Similar distribution of CPAP was detected in centrosomes containing RPCs (Figure 4H). In agreement with their small dimensions, RPC-associated CPAP signal did not exceed the length of ~100 nm and was “embedded” in the mother PCM CPAP signal. Thus, like control nascent PCs, RPCs contain a pool of CPAP available for distancing. Yet, Plk1TD did not associate with the site of RPCs (Figure 4I, left) although Plk1TD associated with MCs even after prolonged Nocodazole treatment (Figure 4I). Thus, without MT walls, PCs could not distance despite Plk1TD-induced expansion of MC-associated Cep192 indicating that, while Cep192 expansion may contribute to distancing in control cells, it is not able to promote distancing of immature PCs.

### Rudimentary procentrioles maintain reduplication block during G2 arrest

In cells arrested in G2 by Cdk1 inhibitor RO-3306 (RO), PCs mature and distance, and MCs reduplicate. These processes are driven by endogenous Plk1 activity (Loncarek et al., 2010). So, we employed this additional reduplication assay to further test the ability of RPCs to distance. Cells were shaken off, re-plated and treated with Nocodazole and Cdk1 inhibitor (Figure 4- figure supplement 2A). Such cells duplicate centrioles, reach G2 phase after ~27 h after shake off, and distance and reduplicate centrioles several hours later. Microscopy analysis showed that, throughout G2 arrest, RPCs remained closely associated with MCs and maintained reduplication block (Figure 4- figure supplement 2B, RO+Noc 8h). The addition of Nocodazole at the beginning of the G2 arrest (27h after shake off) did not prevent PC distancing, showing again that the external MT forces are not necessary for this process. Similarly, inhibition of MT dynein motors by Ciliobrevin allowed initiation, maturation and distancing to occur normally (Figure 4- figure supplement 2B). Washout of Cdk1 inhibitor from G2-arrested cells results in their rapid mitotic entry (Loncarek et al., 2010; Vassilev et al., 2006). In mitotic cells containing RPCs, cartwheel protein STIL was, as expected (Arquint and Nigg, 2014), removed from RPCs (Figure 4- figure supplement C). Thus, as anticipated, RPCs gradually destabilized after mitosis as they lacked MT walls and could not covert to centrosomes (Izquierdo et al., 2014) after mitosis (data not shown).

## Discussion

Mother centrioles and procentrioles are closely associated (engaged (Tsou and Stearns, 2006a)) during interphase, which is a critical aspect contributing to the regulation of centriole number in cycling cells, since the premature loss of such association precedes and leads to the pathological process of centriole reduplication. Nevertheless, by the end of the cell cycle, centrioles lose their close association and by the time of mitotic entry a MC and a PC can be distanced by even >100 nm (Shukla et al., 2015; Vorobjev and Chentsov, 1982). However, surprisingly little is known about the mechanisms that keep centrioles associated in S phase, about the physiological significance of their distancing in late G2, and about how centriole distancing is regulated. Contrary to that, the mitotic events mediated by Separase and Pericentrin resulting in the final separation of two, previously, distanced centrioles, have received more attention and have been better characterized (Kim et al., 2019a; Kim et al., 2015; Lee and Rhee, 2012; Matsuo et al., 2012),

In this work, we elucidated the mechanisms that underly centriole distancing in S phase cells expressing active Plk1, the main promotor of centriole distancing and reduplication. We used nano-scale imaging in combination with biochemical and genetic tools, to dissect the specific role of Pericentrin, Separase, PCM expansion and PC maturation in centriole distancing and reduplication. Our study revealed several novel and important findings. It importantly showed that Separase/Pericentrin remodeling does not occur during centriole distancing and reduplication. This established a clear distinction between the mitotic process of centriole separation and interphase process of centriole distancing that leads to reduplication. While mitotic events involve proteolytic cleavage and post-mitotic reduction of PCM material, interphase events are characterized by accumulation of PCM material to centrosomes. Our nano-scale imaging has revealed that the expression of active Plk1 in S phase cells pushes centrosomes toward a G2-like state, characterized by the accumulation of Cep192 and ɤ-tubulin around MCs. Further, our study demonstrated similarities between PCs in cycling G2 cells and PCs undergoing pathological distancing after Plk1 expression. Using high-resolution imaging, we were able to demonstrate that PCs from cycling cells associated with multiple PCM components already in G2 and prophase, which is when PCs normally distance from their MCs. The same accumulation of PCM components around PCs, although accelerated and amplified, occurs during pathological centriole distancing in interphase cells after Plk1 overexpression. This is a conceptually important finding because human PCs were not known to associate with PCM components before mitosis.

To parse out the effects of MC PCM expansion and PC PCM accumulation on distancing, we inhibited the accumulation of Cep192 (and, in turn, other PCM components) by siRNA. In another set of experiments, we prevented PCs from accumulating PCM components by inhibiting the formation of their microtubule (MT) walls. We found that the absence of PC MT walls, but not the lack of expansion of MC PCM, entirely inhibited centriole distancing. PCs lacking MT walls contained all core centriolar initiating factors, could impose centriole reduplication block within centrosomes and were remarkably stable. But such PCs could not mature and remained adjacent to MCs even if exposed to high Plk1 activity. So, even though the expansion of MC PCM has occurred, it could not drive distancing of immature PCs.

Regardless of the downstream mechanisms which lead to PC distancing, and have yet to be uncovered, our study clearly demonstrates that PC MTs mediate their distancing, controlling their association with MCs, and ultimately controlling centriole reduplication. Based on the striking similarity between physiological and pathological centriole distancing, we propose that centriole reduplication is a consequence of hijacked and amplified PC maturation in response to perturbed cell cycle conditions characterized by unscheduled activity of Plk1 and prolonged cell cycle duration.

## Materials and methods

### Cell culture

HeLa cells constitutively expressing Centrin1-GFP fusion protein (Piel et al., 2000) were used in the study. These cells were additionally engineered to express inducible active Plk1 (Plk1TD, carrying activating T210D mutation) fused with RFP protein, under doxycycline (Dox) inducible promotor (Kong et al., 2014). Cells were grown in DMEM medium (Thermo Fisher Scientific; 10569010) supplemented with 10% Fetal Bovine Serum (FBS) (Omega Scientific; FB-12) at 37 °C in a humified environment with 5% CO2. For microscopy analysis, cells were plated on round 25 mm 1.5-thick high tolerance cover glasses (Warner Instruments; 64-0735). For live-cell imaging, coverslips with cells were mounted in Attofluor Cell chambers (Thermo Fisher Scientific; A7816) in complete CO2-independent medium (Thermo Fisher Scientific; 18045088).

### Cell synchronization and treatments

Cells were synchronized by mitotic shake off. Mitotic cells were shaken off from the tissue culture flasks containing logarithmically growing cells and re-plated on fresh petri dish or on glass coverslip. After shake off cells exit mitosis and reach S within 9-10 h, late G2 within 18-19 h and mitosis within ~21 h after shaking off. To arrest cells in S phase, 2 mM hydroxyurea (HU) (Sigma; H8627) was added 2 h after the shake off. Expression of Plk1TD-RFP was induced with 1 μg/ml Doxycycline (Dox; Sigma-Aldrich). 110 nM of GSK461364 (GSK, Selleckchem; S2193) was used to inhibit Plk1 activity. 10 μM of RO3306 (Tocris; 4181) was used to inhibit Cdk1 activity and arrest cells in G2. MT polymerization was inhibited with 2 μM Nocodazole (Noc; Tocris; 1228). Ciliobrevin (Ciliobr, EMD-Millipore; 250401) was used to inhibit dynein-mediated transport. To generate PCs without MT walls, cells were treated with Nocodazole according to the experimental design detailed in Figure 4- figure supplement 1. To remove Nocodazole from cells and allow centriole MT growth, cells were washed three times for 2 min in pre-warmed complete medium containing HU.

### Identification of cell cycle stages

During microscopy, cell cycle phases were identified as follows: G1 was recognized by the presence of two single C1-GFP signals and/or the presence of the midbody between two sister cells. Early G1 was recognized by the proximity of the sister cells and their round shape. In S phase, cells contained duplicated centrioles (four C1-GFP signals) and closely positioned centrosomes. G2 was identified by two separated and duplicated centrioles with developed PCs. Prophase was identified by DNA condensation. In some experiments, cells were co-labeled with the DNA duplication marker PCNA which is present in the nuclei of late S phase cells and absent from nuclei of G2 (Kennedy et al., 2000; Leonhardt et al., 2000), to distinguish late S from early G2 cells. Stages of mitosis were recognized by characteristic DNA morphology.

### Protein depletion by siRNA

The following siRNA oligonucleotides were used: Separase siRNA; 5’-GCAGGUUCUGUUCUUGCUUGA-3’, Cep 192; 5’-GCUAGUAUGUCUGAUACUUGG-3’, and siRNA targeting Luciferase; 5’-UAAGGCUAUGAAGAGAUAC-3’ (Dharmacon) as a control. Cells, growing on glass coverslips or in 75 cm^2^ cell culture flasks, were transfected 2h after shake off with 300 nM of siRNA oligonucleotides and Oligofectamine (Thermo Fisher Scientific; 12252011). Cells were fixed for immunofluorescence, lysed for immunoblot analysis or shaken off and re-plated, as described for individual experiments in the main text and in Figure 2-figure supplement 1.

### Expression of HA-Separase

To analyze distancing and re-duplication after overexpression of Separase and Plk1TD, cells were synchronized by mitotic shake off and re-plated, arrested with HU 2 h later, and transfected at 4-6 h with a plasmid encoding HA-Separase (pCS2+HA-hSeparase was a gift from Marc Kirschner, Addgene plasmid # 33018), Plk1TD expression was induced 8-10 h after shake off and cells were fixed at 28 h to analyze distancing or at 32 h (20 h of S phase arrest) to analyze reduplication. To evaluate centriole distancing, samples were analyzed by SIM imaging, as described below.

### Immunolabeling

Cells were fixed with 1.5% formaldehyde in PBS for 4-6 min and post-fixed with ice cold methanol for 5 min at −20°C. After several washes in PBS, the cells were blocked for 15-20 min in PBS containing 1% BSA and 0.05% Tween-20. To detect centrosomal Separase signal, cytosolic extraction was performed with 0.1% Triton in PHEM buffer for 45 s, followed by fixation with 2% formaldehyde and several washes with PBS before blocking (Yuan et al., 2009). After blocking, cells were incubated with primary antibodies diluted in blocking buffer for 1-3 h at 37°C or overnight at 4°C. After washing in PBS, cells were exposed to secondary antibody diluted in blocking buffer for 1 at 37°C. DNA was stained with Hoechst 33342 (Thermo Fisher Scientific; H3570).

Secondary antibodies Alexa Fluor 488 anti-mouse (Thermo Fisher Scientific; A11029), Alexa Fluor 488 anti-rabbit (Thermo Fisher Scientific; A11034), Alexa Fluor 555 anti-mouse (Thermo Fisher Scientific; A28180), Alexa Fluor 555 anti-rabbit (Thermo Fisher Scientific; A21429), CF647 anti-mouse (Biotium; 20042), and CF647 anti-rabbit (Biotium; 20045) were used at a 1:800 dilution to label primary antibodies for 24 h. The following primary antibodies were used for immunofluorescence: mouse anti-SAS-6 (Santa Cruz; sc-81431) at 1:250, rabbit anti-STIL (A gift from Dr. Andrew Holland) at 1:500, rabbit anti-Pericentrin (Abcam; ab4448) at 1:4000, rabbit anti-Cep135 (Bethyl; A302-250A) at 1:1000, goat anti-CPAP N-term (Santa Cruz; sc-66747) at 1:500 and rabbit anti-CPAP C-term (Proteintech; 11577-1AP) at 1:500, rabbit anti-Cep63 (Millipore; 06-1292) at 1:1000, mouse anti-γ-tubulin (Sigma; T6557) at 1:5000, rabbit anti-ɤ-Tubulin (Sigma; T3559) at 1:500, rabbit anti-Cep192 (Bethyl; A302-324AM) at 1:500, mouse anti-HA (Santa Cruz; sc-7392) at 1:250, rabbit anti-HA (Proteintech; 51064-2-AP) 1:250, Rabbit anti-Cep152 (Bethyl; A302-479A) at 1:3000, mouse anti-Plk1 (Santa Cruz; sc-81431) at 1:500, and mouse anti-Separase (Santa Cruz; sc-390314) at 1:300. To stain PCNA, a prelabeled rabbit anti-PCNA (Proteintech; 10205-2-AP) was used at 1:200. PCNA antibody was pre-labeled using a Mix-n-StainCF-568 Dye Antibody Labeling Kit (Biotium; 92275) accordingly to the manufacturer protocol.

### Widefield microscopy and Structured Illumination Microscopy (SIM)

Widefield imaging was performed with an inverted Eclipse Ti microscope (Nikon), equipped with OrcaFlash4 camera (Hamamatsu), Intensilight C-HGFIE illuminator, 100x NA 1.45 Plan Apo objective, using 1.5x magnifying tube lens. 200 nm thick Z-sections spanning the entire centriole or central centriole region were acquired, as needed. SIM was performed using N-SIM (Nikon), equipped with 405, 488, 561, and 640 nm excitation lasers, Apo TIRF 100x NA 1.49 Plan Apo oil objective, and back-illuminated 16 μm pixel EMCCD camera (Andor; DU897). Hundred nm-thick Z sections were acquired in 3D SIM mode and reconstructed to generate a final image using Nikon NIS-Elements software.

### Stochastic optical reconstruction microscopy (STORM)

Before STORM imaging, coverslips with immunolabeled cells were layered with 100 nm tetra-spectral fluorescent microspheres (Thermo Fisher Scientific; T7279), which served as fiducial markers. Coverslips were mounted in Attofluor Cell chambers (Thermo Fisher; A7816), filled with STORM buffer (25 mM β-mercaptoethylamine [Sigma; 30070], 0.5 mg/ml glucose oxidase [Sigma; G2133], 67 μg/ml catalase [Sigma; C40], 10% dextrose [Sigma; D9434], in 100 mM Tris, pH 8.0) and covered with an empty coverslip. 3D STORM imaging was performed on Nikon N-STORM4.0 system using Eclipse Ti2 inverted microscope, Apo TIRF 100X SA NA 1.49 Plan Apo oil objective, 405, 561 and 647 nm excitation laser launch and a back-illuminated EMCCD camera (Andor; DU897). The 647 nm laser line (~150 mW out of the fiber and ~90 mW before the objective lens) was used to promote fluorophore blinking. The 405 nm laser was used to reactivate fluorophores. The 561 nm laser was used to record the signals of fiducial markers. 20,000 to 30,000 time points were acquired every 20 ms at a 50 Hz frame rate. NIS Elements (Nikon) was used to analyze the data. Prior to STORM imaging the position of the Centrin1-GFP-labeled centrioles and of the CF647-labeled target protein was recorded in widefield mode. Multiple STORM images were taken from two or more independent experiments. The resolution in our STORM experiments was estimated based on the analysis of cytoplasmic MTs, as described in (Bowler et al., 2019).

### Analysis and presentation of STORM data

A rainbow Z-color coding scheme, which typically spanned 650 nm of a working Z-imaging range, was used for signal presentation. The signals closer to the coverslip were presented in red and those further from the coverslip in blue. 3D STORM data is presented as a projection of the entire 3D volume containing centrosomal signal. Sometimes only the central portion of the STORM signal is shown for clarity purposes. 3D STORM volume viewer was used to analyze data in 3D. An outline, based on the 3D analysis, was drawn for each presented centriole, illustrating approximate centriole’s position, orientation, and length. The outline was proportionally scaled for each panel to reflect known centriole dimensions which are: ~230 nm for MC diameter; ~200 nm for control PC diameter, and ~160 and 100 nm for the diameter and length of RPCs. Widefield images of centrioles analyzing by STORM were scaled 3x with bicubic interpolation.

### Electron microscopy

For correlative live and electron microscopy analysis, cells growing on the coverslip were assembled in an Attofluor Cell chambers and imaged by Nikon Eclipse Ti inverted microscope using 100x NA 1.45 Plan Apo objective, Yokogawa spinning disc, 405, 488, 561 and 640 nm laser launch (Agilent technology MCL-400), back-illuminated EMCCD camera (Andor, DU888), and a 2x relay lens. A Nikon DS-U3 camera was used to record DIC images. After live cell recording cells were fixed in 2.5% glutaraldehyde (Sigma; G5882) and 0.25% formaldehyde in 1× PBS (pH 7.4) for 1 h at RT. Centriole position within the target cells was recorded and the position of the cells was marked on the coverslip. Cells were pre-stained with 1% osmium tetroxide (Electron Microscopy Sciences; 19100) and 1% uranyl acetate (Electron Microscopy Sciences; 22400), dehydrated in ethanol, and embedded in EMbed-812 resin (Electron Microscopy Sciences; 13940). 80 nm thick serial sections were sectioned, placed on the formvar-coated copper grids (SPI Supplies; 2330P-XA), stained with uranyl acetate and lead citrate, and imaged using a transmission electron microscope (Hitachi) or FEI Spirit transmission electron microscope operating at 80 kV. The alignment of the serial sections and image analysis and was performed in Photoshop and Fiji.

### Measurement of Centrin-GFP distances

To measure distance between MCs and PCs, images of centrioles were acquired (using SIM) and the X Y and Z coordinates of Centrin1-GFP centroids were determined from the recordings. The distance between two signals was then calculated using Pythagorean theorem in three dimensions. Centrioles were also considered distanced, when they were visibly associated with pericentriolar components or Plk1, as such centrioles showed an increased distance from MCs if analyzed by STORM.

### Immunoblotting

Cells were lysed in reducing Laemmli buffer (100mM M Tris-HCl pH 6.8, 20% glycerol, 4% SDS, 200mM DTT, 0.01% bromophenol blue) or, for quantification of protein concentration, Triton-Lysis buffer (50 mM Tris-HCl pH 7.5, 150 mM NaCl, 1% Triton, 1 mM EDTA, 1 mM PMSF) supplemented with protease cocktail inhibitor (Roche; 11836153001). Proteins were separated by SDS PAGE using 6% gels to resolve PCNT or separase, or 8% gels for lower molecular weight proteins, transferred to a PVDF membrane, blocked in 6% skim milk diluted in diluted in TBS-Tween 0.1% buffer (TBS-T) and incubated with primary antibody in 3% milk in TBS-T for 1 h at RT or overnight at 4°C. Primary antibodies were used as follows: mouse anti-Plk1 (Santa Cruz; sc-17783) at 1:500, rabbit anti-Pericentrin (Abcam; ab4448) at 1:2000, rabbit anti-Separase (Bethyl; A302-215A) at 1:1000, mouse Separase (Santa Cruz; sc-390314) at 1:500, rabbit anti-Lamin A/C (Proteintech; 10298-1-AP) at 1:10,000. HRP-conjugated secondary anti-mouse antibody (Amersham; NA931S) and anti-rabbit antibody (Amersham; NA934VS) at 1:10,000 and Clarity western ECL substrate detection kit (Bio-Rad; 170-5060) were used to detect the signals. Precision Plus (Bio-Rad; 161-0374) was used as protein size marker.

### Statistical Analysis

Statistical difference between two sets of data was determined in Excel using an unpaired, two-tailed Student’s t-test. NS, nonsignificant; P ≤ 0.05: *, P ≤ 0.01: **, P ≤ 0.001: ***. Box-and-whisker plots show the minimum, median line, upper and lower quartile, maximum, and inner and outer points. Histograms show average values ± standard deviation. Sample sizes (the number of counted or measured centrioles and experiment numbers) are indicated on the plots or in the figure legends.

## Acknowledgements

This research was supported by the Intramural Research Program of the National Institutes of Health (NIH), National Cancer Institute to JL.

The authors declare no competing financial interests.

## Author contributions

AVL, conducted most experiments, analyzed the data and prepared the manuscript. DK prepared EM samples. AS conducted some experiments on RCs. CS and KL conducted STORM. JL supervised the project and conducted some microscopy and data analysis. JL and AVL wrote the manuscript. All authors discussed the data and reviewed the manuscript.

## List of supplementary files

**Supplementary File 1.** A table listing all antibodies used in microscopy analysis and detailing their epitopes.

**Table 1.** A list of the antibodies used for Immunolabeling and microscopy analysis. Immunolabeling conditions are described in Materials and Methods section.

| **Protein (human):** | **Protein Length (aa):** | **Targeted Domain:** | **Epitope (aa):** | **Antibody: (source, manufacturer; Cat. No.)** |
| --- | --- | --- | --- | --- |
| Sas-6 | 657 | C-terminus | 404-657 | Mouse, Santa Cruz; sc-81431 |
| STIL | 1288 | C-terminus | 1271-1278 | Rabbit, Gift from Dr. A. Holland |
| Cep192 | 2537 | C-terminus | 1800-1850 | Rabbit, Bethyl; A302-24AM |
| Cep152 | 1710 | N-terminus | 50-100 | Rabbit, Bethyl; A302-479A |
| Cep63 | 703 | Full protein | Full protein | Rabbit, Millipore; 06-1292 |
| Pericentrin | 3336 | N-terminus | 100-600 | Rabbit, Abcam; ab4448 |
| γ-tubulin | 451 | N-terminus | 38-53 | Mouse, SIGMA; T6557 |
| γ-tubulin | 451 | N-terminus | 38-53 | Rabbit, SIGMA; T3559 |
| CPAP | 1338 | C-terminus | 1033-1338 | Rabbit, Proteintech; 11577-1AP |
| Cep135 | 1140 | C-terminus | 900-950 | Rabbit, Bethyl; A302-250A |
| Plk1 | 603 | Middle region | 261-412 | Mouse, Santa Cruz; sc-81431 |
| Separase | 2120 | C-terminus | 1496-1795 | Mouse, Santa Cruz; sc-390314 |
| PCNA | 261 | Full protein | 8-256 | Rabbit, Proteintech; 10205-2-AP |
| HA tag | 9 | - | 1-9 | Rabbit, Proteintech; 51064-2-AP |
| HA tag | 9 | - | 1-9 | Mouse, Santa Cruz; sc-7392 |

**Supplementary File 2.**
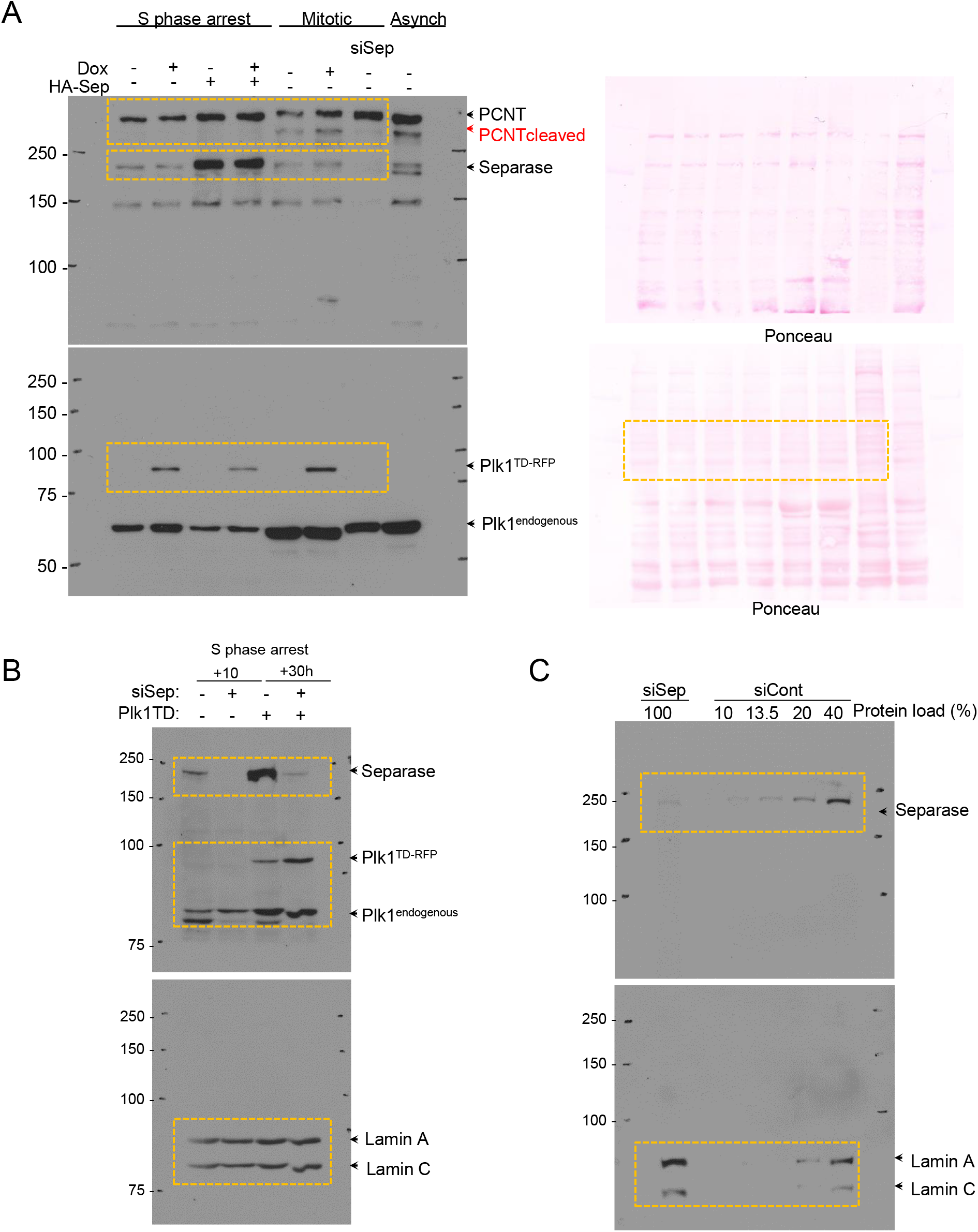
(A) Scans of immunoblots and immunoblotting membranes used in Figure 2A. After immunoblotting, membranes were stained with Ponceau to ascertain the levels of proteins. (B, C) Scans of immunoblots used in Figure 2G. Signals of Lamin A and C were used as a loading control.

**Supplementary File 3.**
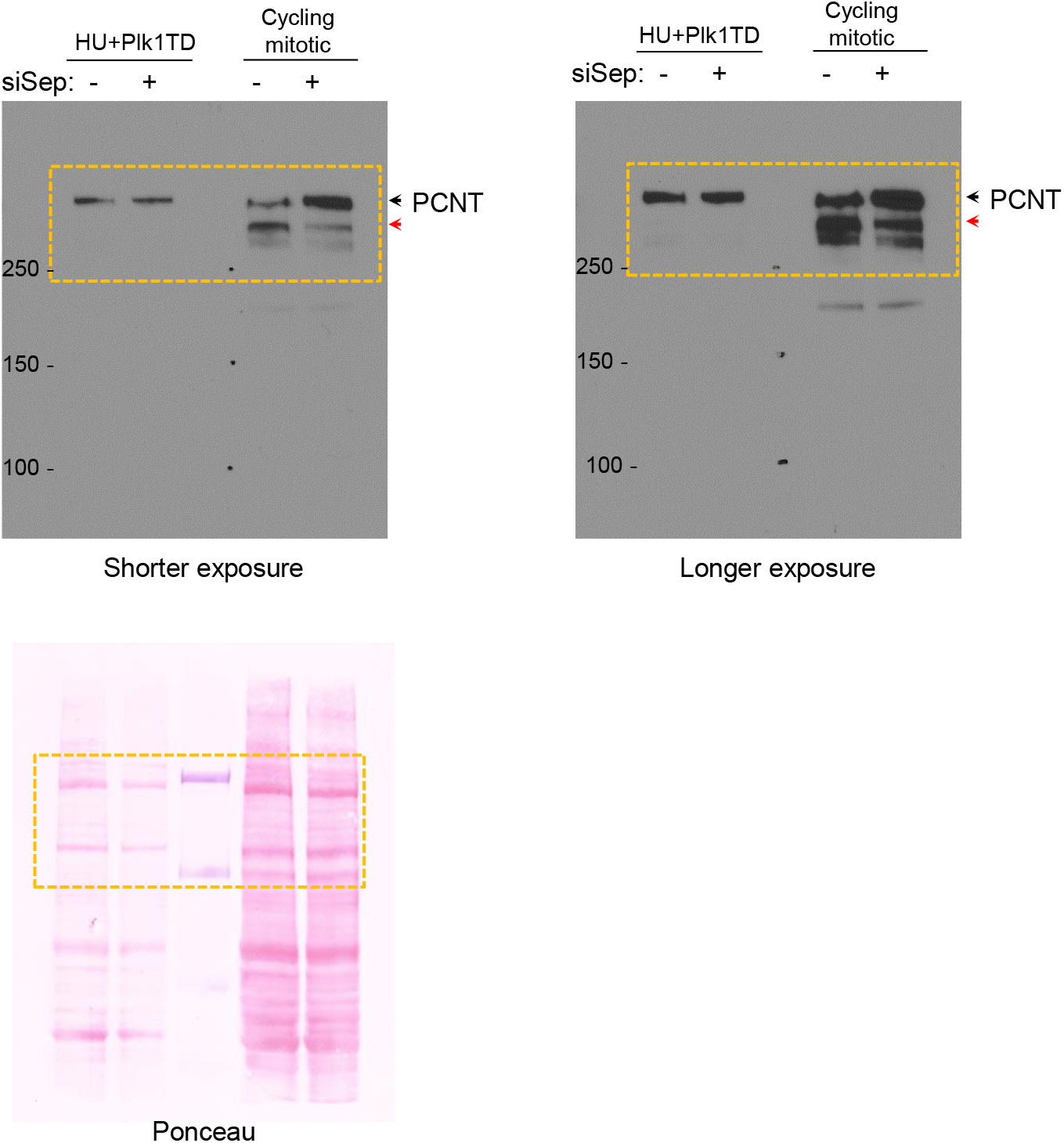
Scans of immunoblot films and membrane shown in Figure 2- figure supplement 1B after short and long exposure. After immunoblotting, membranes were stained with Ponceau to ascertain the levels of proteins.

**Figure 1- figure supplement 1.**
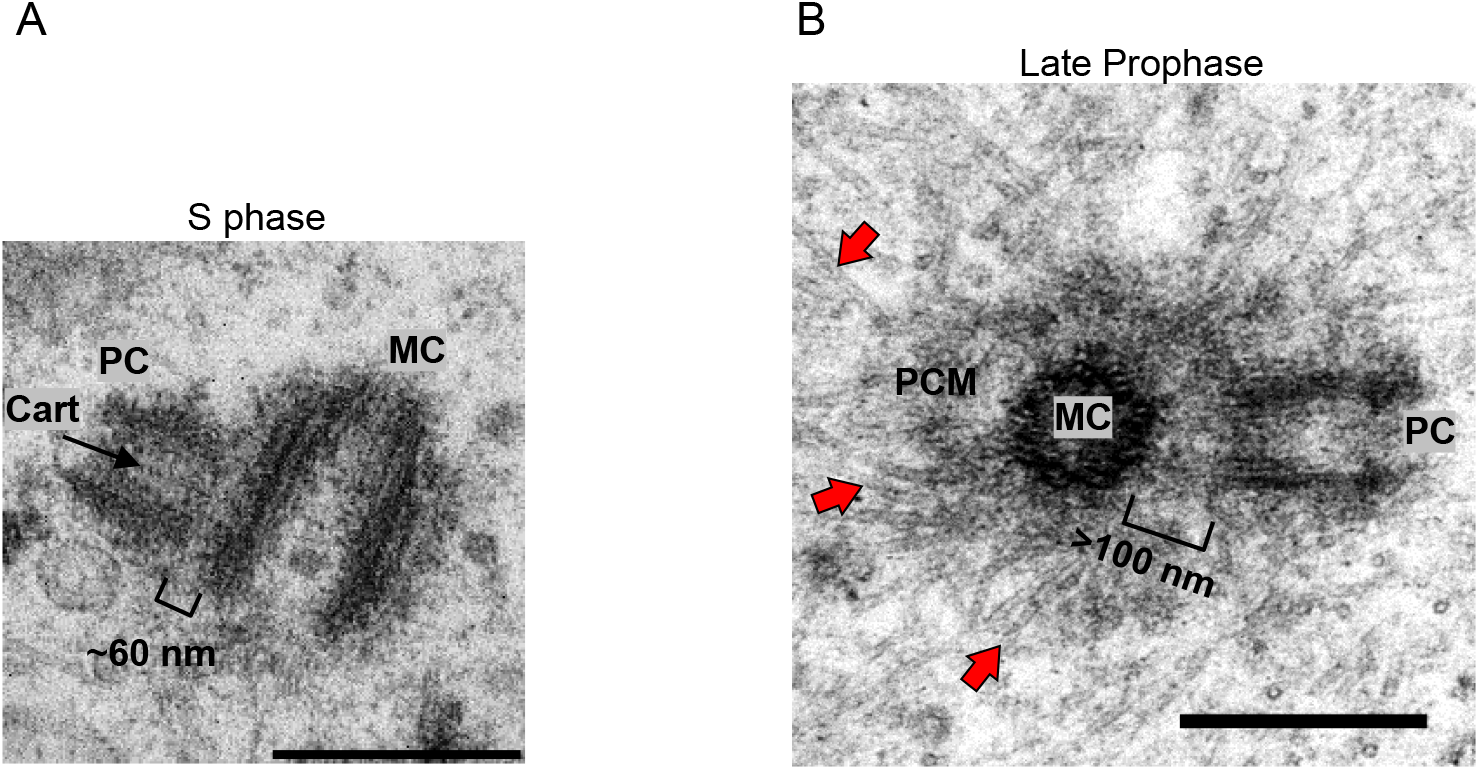
Electron micrographs of centrosomes from S phase and prophase HeLa cells. (A) Duplicated mother centriole (MC) is associated with procentriole (PC), which has formed orthogonally to MC’s microtubule walls. The distance between MC and PC is ~60 nm. Cartwheel components are visible in PC’s lumen (Cart). (B) MC-associated PCM expands in prophase (dark-gray material around MCs, please compare with Figure 1H) and PC is at a larger distance from MC than in S phase cell. Microtubules nucleated by expanded PCM are visible around PCM (red arrows). Scale bars: 0.4 μm.

**Figure 1- figure supplement 2.**
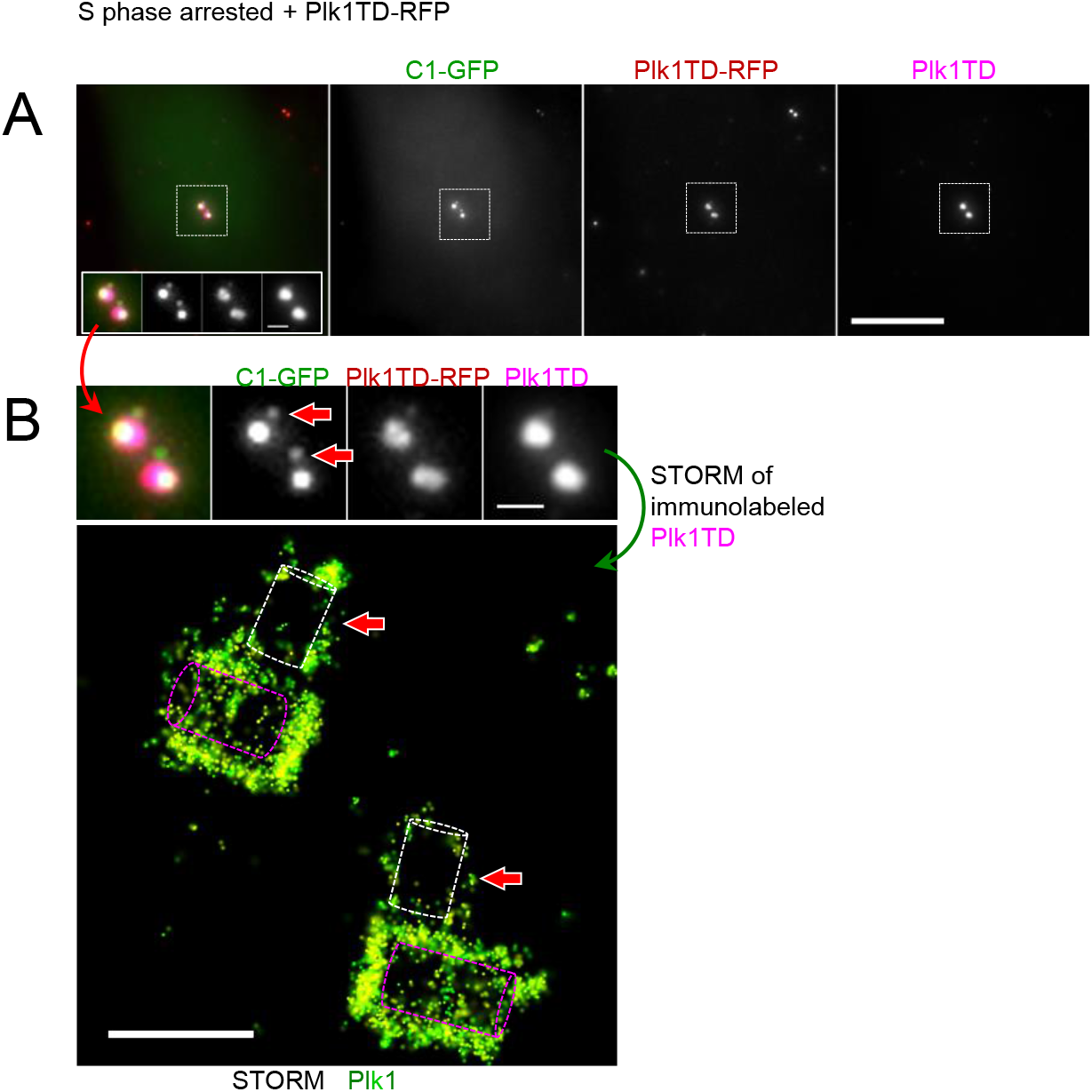
Correlative widefield/STORM analysis of centrosomal Plk1. Cell constitutively expresses C1-GFP to mark centrioles. Plk1TD-RFP (red) is induced by Doxycycline. Cells are fixed and immunolabeled using primary anti-Plk1 antibody and secondary blinking antibody. Please note that immunolabeling detects both, endogenous Plk1 and Plk1TD, although in S phase cells, Plk1TD is predominant after induction (Figure 1A). (B) STORM analysis of Plk1 from (A). Plk1TD is accumulated at low levels on procentrioles (red arrows), which are still associated with mother centriole and orthogonally oriented. A middle section through STORM signal is shown. Purple outlines: mother centrioles, white outlines: procentrioles Scale bars: 10 μm in A, and 1 μm in centriole inserts in A and B, 0.5 μm for STORM in B.

**Figure 2- figure supplement 1.**
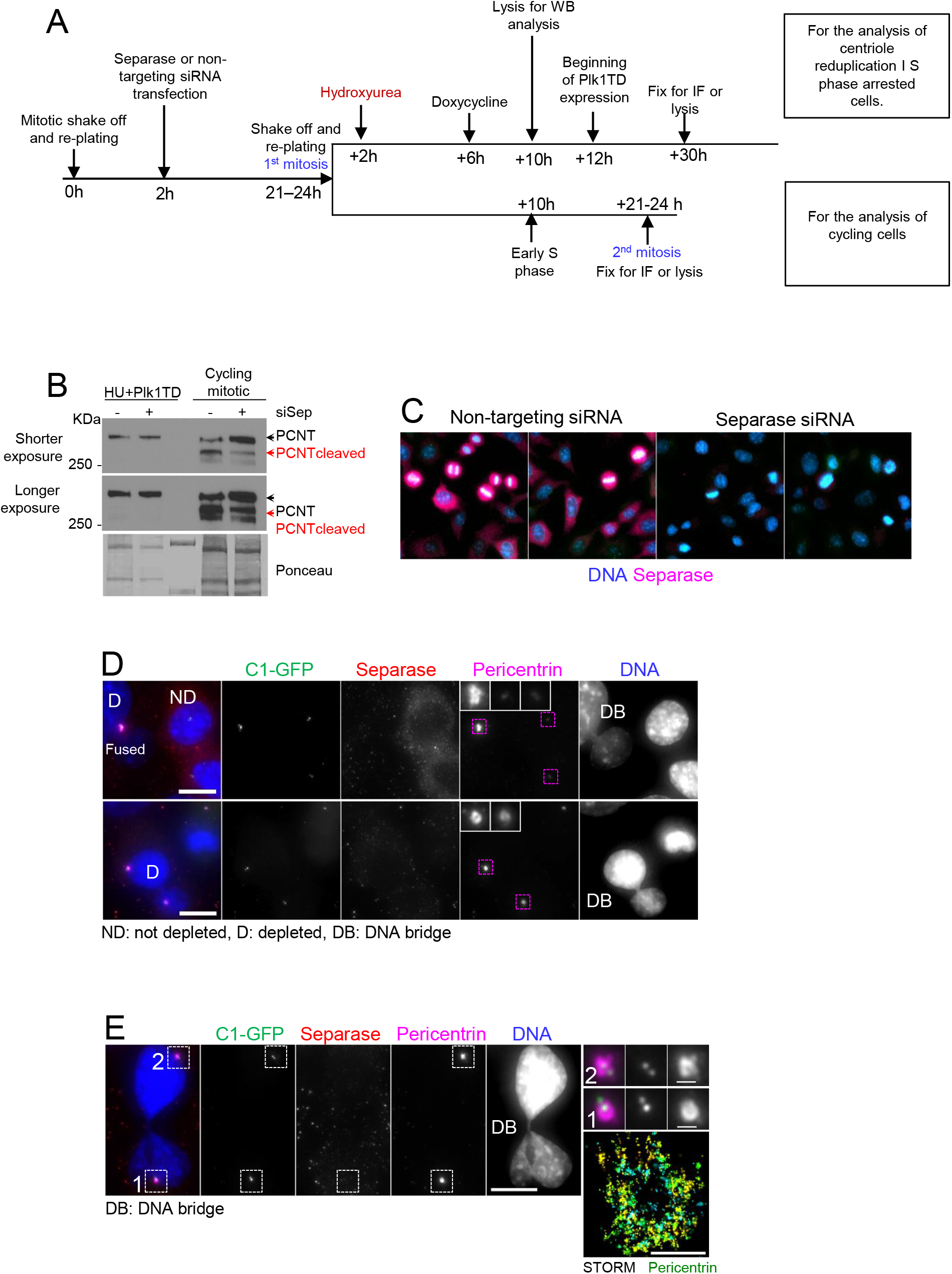
Depletion of Separase in HeLa cells. (A) A scheme depicting the strategy used to generate Separase-depleted S phase arrested and cycling cells. (B) Immunoblot analysis of Pericentrin cleavage after Separase depletion. Please note reduced cleavage of Pericentrin in mitotic Separase-depleted cells. The signal after longer exposure is included to ascertain the lack of Pericentrin cleavage in S phase arrested cells by Hydroxyurea (HU) after Plk1TD expression. (C) Immunofluorescence analysis of cell population transfected with non-targeting or Separase siRNA. Separase labeling (magenta) is significantly reduced in the population of cycling cells. DNA is labeled in blue. (D) Centrosome-associated Pericentrin levels are higher in Separase-depleted postmitotic cells in comparison with cells with detectable Separase (ND cells). DNA bridges connecting two sister cells (DB) are the hallmark of Separase-depleted cells and are visible in cells lacking Separase. (E) Two sister cells from cell culture depleted for Separase associated with DNA bridge. Pericentrin is still abundant on centrosomes. STORM analysis of centrosomes #1, shows its distribution around centrioles. Scale bars: 10 μm and 1μm for for centriole inserts.

**Figure 2- figure supplement 2.**
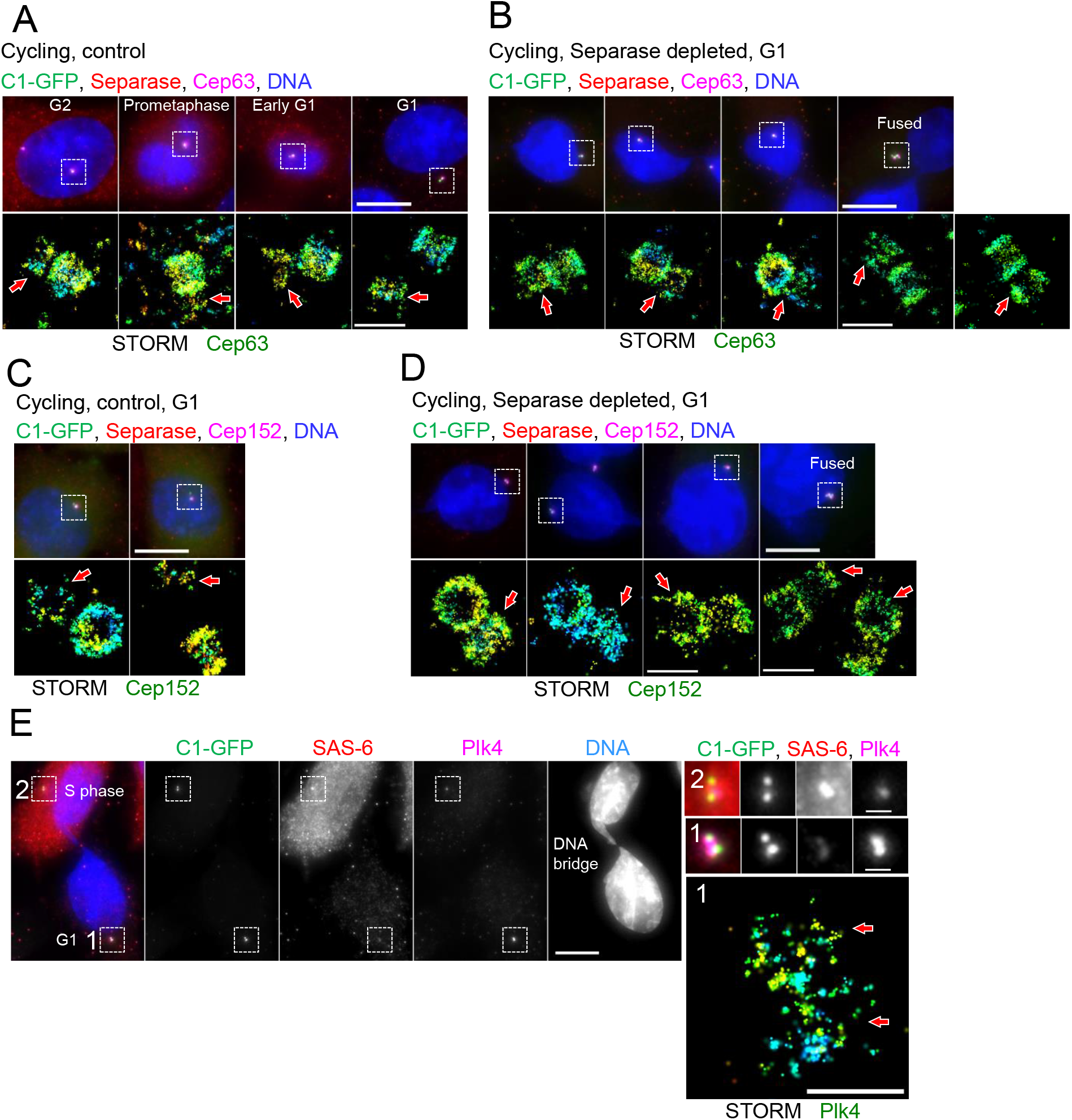
Localization of Cep63, Cep152 and Plk4 in Separase-depleted HeLa cells. Correlative widefield/STORM analysis is performed in all panels. Centrosomes in white squares were analyzed by STORM. Red arrows point to a younger centriole from a pair. (A) Centrosome-associated Cep63 in cycling cells in various stages of the cell cycle. (B) Centrosome-associated Cep63 in Separase-depleted cycling G1 cells. (C) Centrosome-associated Cep152 in cycling G1 Cells. (D) Centrosome-associated Cep152 in Separase-depleted cycling G1 cells. (E) Two sister cells from cell culture depleted for Separase connected with a DNA bridge. Centriole initiator factor SAS-6 is variably expressed in two sister cells and present on centrosomes of two cells at different levels. Plk4, centriole initiating kinase, is variably present on centrosomes. STORM analysis of centrosome 1, shows its distribution around centrioles. Scale bars: 10 μm and 1 for centriole inserts.

**Figure 3- figure supplement 1.**
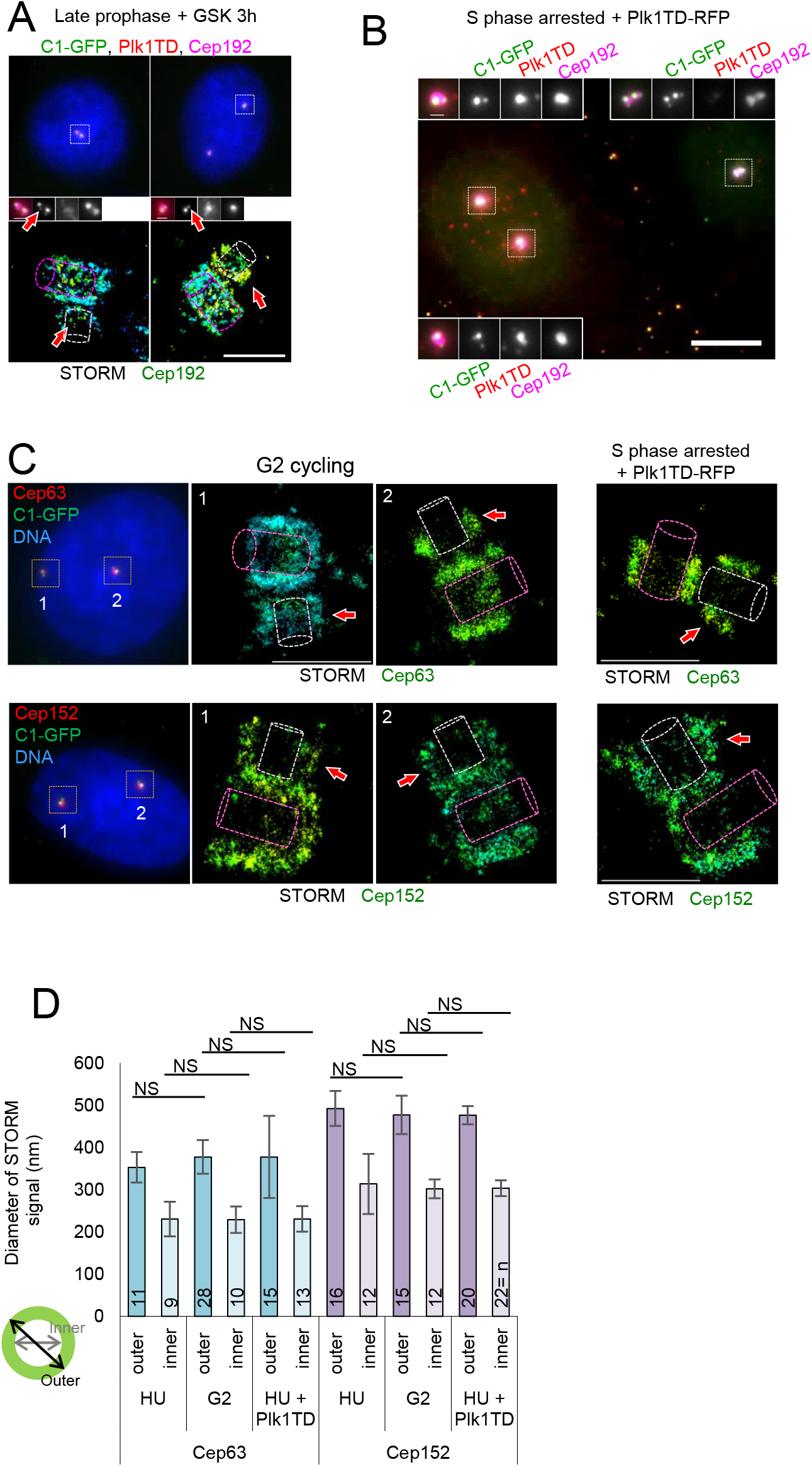
STORM analysis of PCM components in cycling and S phase arrested cells expressing Plk1TD. (A) Correlative widefield and STORM analysis of Cep192 localization in prophase HeLa cells treated with Plk1 inhibitor GSK461364 for 3 h. Plk1 inhibition prevents mitotic expansion of Cep192. Arrows point to Cep192 signal associated with procentrioles. (B) Two S phase arrested cells with different levels of Plk1 TD are shown. Centrosomes with high levels of Plk1 TD are associated with higher levels of PCM component Cep192. (C) Correlative wide-field and STORM analysis of Cep63 and Cep152 in cycling G2 cells and S phase arrested HeLa cells expressing Plk1TD. Red arrows point to procentrioles associated with Cep63 and Cep152 signal. (D) Quantification of Cep63 and Cep152 STORM signals. Inner and outer diameters were measured as illustrated. Histograms show an average ± SD from three independent experiments. Purple outlines: mother centrioles, white outlines: procentrioles. The sample size (n) is shown on the graphs. ns: nonsignificant. Scale bars: 0.5 μm for STORM in all panels, 1 μm for centriole inserts, 10 μm in B.

**Figure 4- figure supplement 1.**
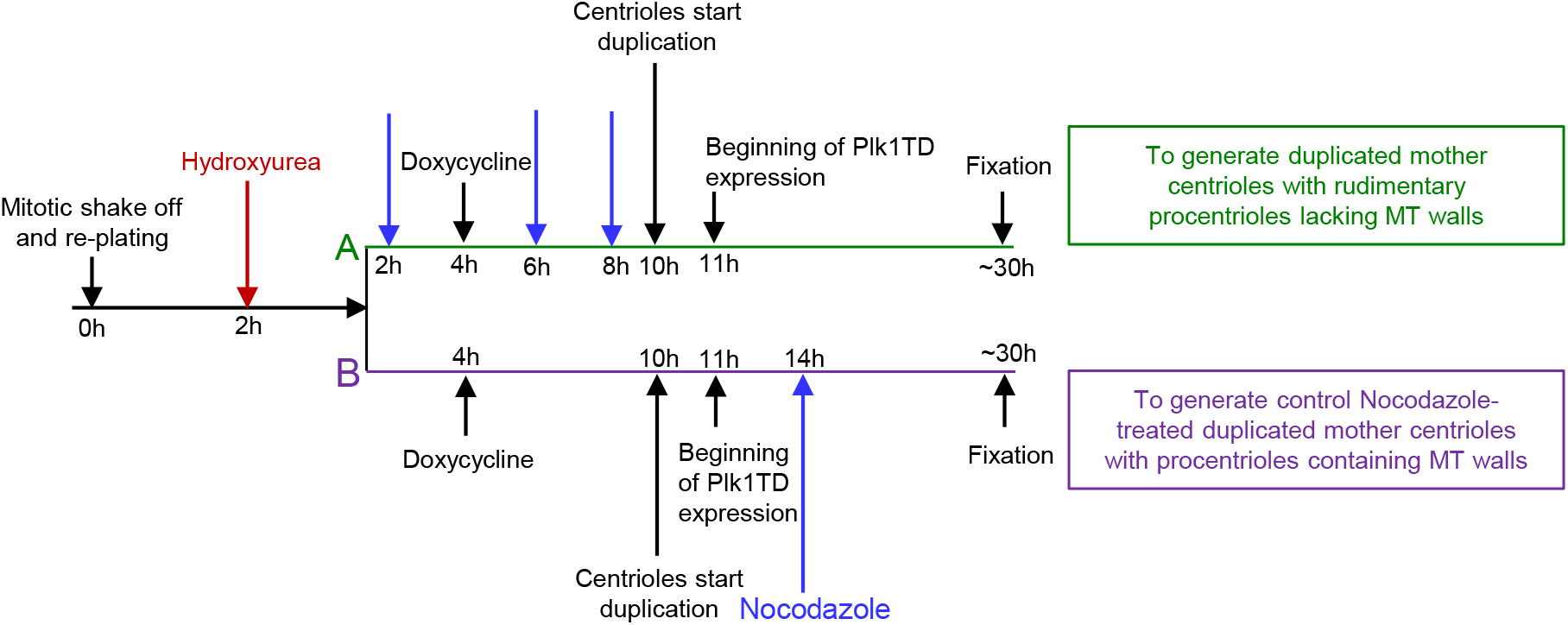
A scheme illustrating experimental design used to generate procentrioles lacking microtubules. To generate duplicated mother centrioles containing procentrioles lacking microtubules, cells were treated with Nocodazole at various points in G1 (A). To generate control Nocodazole-treated duplicated mother centrioles containing short procentrioles with microtubules, Nocodaloze was added after centrioles had duplicated (B). Plk1TD: active form of Plk1, MT: microtubule. Doxycycline was used to induce Plk1TD expression.

**Figure 4- figure supplement 2.**
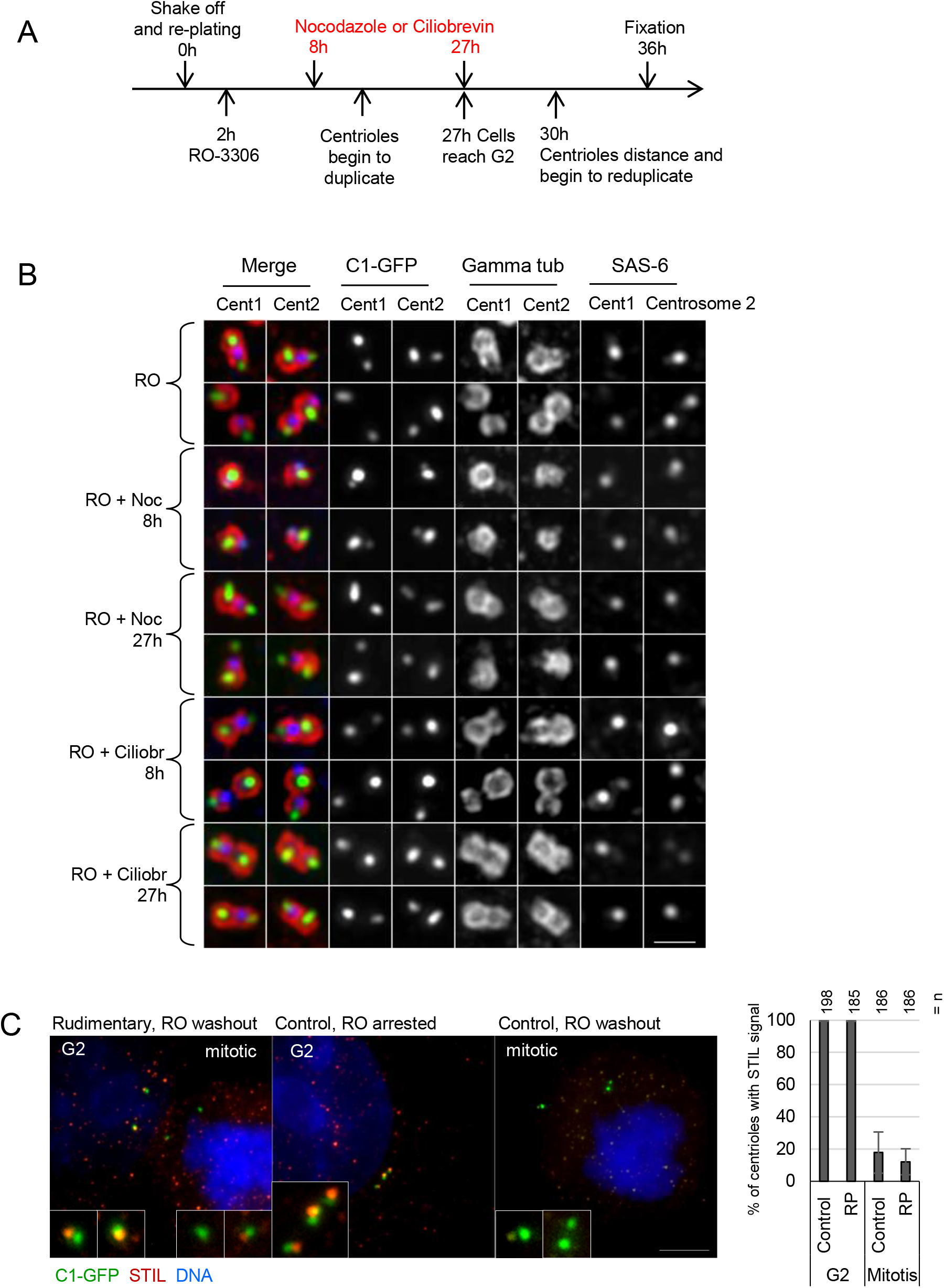
Non-centriolar microtubules are dispensable for centriole distancing and reduplication in interphase. (A) Scheme illustrating experimental design used to test the ability of procentrioles lacking microtubule (MT) walls to distance and allow reduplication of their mother centrioles. Nocodazole (Noc) was added in G1 cells to prevent microtubule incorporation to nascent procentrioles. Ciliobrevin (Ciliobr) was added to prevent dynein-mediated transport. RO-3306 (RO) is a Cdk1 inhibitor used to arrest cells in G2. (B) The analysis of G2 arrested cells treated as explained in (A) for their ability to mature and distance centrioles in the absence of microtubules (Noc) and MT-dependent transport (Ciliobr). In cells treated with RO only, procentrioles (weaker C1-GFP signals) accumulate PCM component gamma tubulin and distance from mother centrioles. SAS-6 signal marks the proximal ends of procentrioles. In cells treated with Nocodazole from G1, procentrioles remain in the vicinity of mother centrioles. Addition of Nocodazole from early S, after procentriole MT walls had assembled, does not prevent accumulation of PCM around procentrioles and distancing. Ciliobrevin does not inhibit accumulation of gamma tubulin by procentrioles or their distancing. (C) Microscopy analysis of cartwheel protein STIL in centrosomes containing rudimentary procentrioles or control procentrioles. STIL associates with procentrioles in G2 arrested cells but is removed upon mitotic entry induced by RO washout. Right: Quantification of cells with centrosomal STIL signals. Histogram shows an average ± SD from three independent experiments. The sample size (n) is shown above the graph. Scale bar: 1 μm.

